# Joint Moment–Angle/Velocity Relations in the Hip, Knee, and Ankle: A Meta-Visualization of Datasets

**DOI:** 10.1101/2024.06.22.600197

**Authors:** Ziyu Chen, David W. Franklin

## Abstract

Joint moment is a prominent kinetic property in biomechanical investigations, whose pattern and magnitude unravel the characteristics of musculoskeletal motion and muscle biomechanics. Nonetheless, the relations of joint moment with joint angle and velocity are complicated, and it is often unclear how the kinetic capacity of each joint varies in different configurations. With common techniques in systematic review, we collected a total of 985 passive, isometric and isokinetic joint moment datasets from literature and visualized the major joint moment–angle and moment–velocity relations in the hip, knee, and angle. The findings contribute to the understanding and analysis of musculoskeletal mechanics, as well as providing reference regarding the experimental design for future moment measurement.

## 1. Introduction

The moment of force in the joint is a major interest in the study of human movement, as it directly associates kinematics with muscle biomechanics (Anderson and Pandy, 1999; Franklin et al., 2003; Hu et al., 2020; Zajac, 1993). The pattern and magnitude of joint moments, calculated via inverse kinematics, is often used in biomechanical analysis as a substitute for the timing and intensity of muscle activities (Franklin et al., 2003; Milner and Franklin, 2005; Raffalt et al., 2016; Robertson et al., 2008), which cannot easily be obtained during dynamic motions (De Luca, 1997). Similarly, maximal joint moment at different joint angles and rotational velocities is an important indicator of muscle strength, and is often correlated with movement performance or dysfunction (Beynnon et al., 2001; Boling et al., 2009; Jones et al., 2009; Tsiokanos et al., 2002). Data such as isometric and isokinetic moments provide valuable references for medical diagnosis (Ekdahl and Broman, 1992; O’Neill et al., 2019), rehabilitation (Perhonen et al., 1992; Shaffer et al., 2000), and athletic training (Daneshjoo et al., 2013; Handel et al., 1997), as well as play an essential role in the calibration of musculoskeletal models (Anderson and Pandy, 1999; Delp et al., 1990; Holzbaur et al., 2005), which are a convenient tool for the estimation of muscle force or activation (Delp et al., 2007).

Theoretically, joint moment is not particularly difficult to measure: When the actuated limb enacts against an object, a contact force is in effect, and joint moment can be calculated as the product of this force and the distance between the contact point and rotation center. However, despite the importance of isometric and isokinetic moment data or the simplicity of moment measurement, it is still hard to find constructive datasets that clearly depict the relations of moment–angle/velocity in each joint (Anderson et al., 2007; Silder et al., 2007). There is a fundamental reason behind this reality. Unlike the muscle force–length relation, the joint moment–angle relation is not necessarily one-dimensional (Chen et al., 2024; Prilutsky and Zatsiorsky, 2002). For any monoarticular muscle, there are theoretically three degrees of freedom (DoF) that this muscle actuates. In the hip, for instance, all of its muscles should have some level of contribution to its extension/flexion, abduction/adduction, and external/internal rotation. Reversely, a movement in any one of the DoFs affects the lengths of all relevant muscles, their maximal force outputs, and ultimately the maximal moment outputs in all directions. For example, the maximal hip abduction moment will change not only when the hip abducts, but also when the hip flexes or rotates. Therefore, the relation of hip abduction moment with hip position is theoretically three dimensional, and it is the same for hip moments in the other five directions.

This problem is further complicated in biarticular muscles such as the rectus femoris and hamstrings; that is, position of the knee could potentially affect hip moments, and vice versa (Herzog and ter Keurs, 1988; Savelberg and Meijer, 2003). As a result, to fully understand the moment–angle relation of a joint, data must be obtained from a combination of various joint positions involving all related DoFs, which is a time-consuming task. In practice, researchers tend to focus on the sagittal plane, e.g., by keeping the hip in the neutral position of abduction/adduction and external/internal rotation to study the relation between hip extension (or hip flexion, knee extension, knee flexion) moment and the flexion angle of either the hip or knee. Even still, any of these moment–angle relations is two dimensional, and the number of joint positions for isometric measurement increases quadratically compared to the case that would exist if the relation was only one dimensional.

Furthermore, how fast the joint rotates also has an impact on its moment output, and this impact may vary with joint angle (Anderson et al., 2007; Khalaf et al., 2000; Hussain and Frey-Law, 2016). In other words, this relation of joint moment–velocity should always contain one more dimension than the accordant moment–angle relation. Fortunately, the measurement of an isokinetic moment is no more complicated than that of an isometric moment, because in a single isokinetic test, the joint needs to move over a specific range covering joint angles measured in multiple isometric tests. However, it is very common for researchers to measure the isokinetic moment over some range but take the peak or mean value in representation of the moment at the test angular velocity (Boling et al., 2009; Gajdosik, 2002; Vandervoort et al., 1990). By doing so, the muscle force–velocity relation is assumed to be constant across all muscle lengths, which is not necessarily the case.

As a result, there is still great uncertainty behind the seemingly simple question of how much moment a joint could generate in a specific position and rotational velocity. To reveal such potentially complicated biomechanical relations, more measurements are of course required to cover a wider range of joint configurations. Nevertheless, before designing a series of large experiments, it is necessary to find out which parts of the joint space are already well-studied and which parts require further measurements.

Here, we extensively collect existing datasets from literature constituting joint moment–angle and joint moment–angular velocity relations for six DoFs in the lower limb. We approach the issue using common techniques in systematic review and we aim to visualize the data with sufficient details to provide reference values for biomechanical analysis, especially musculoskeletal modeling.

## 2. Methods

In this study, we focus on six major DoFs in the lower limb:

1. hip extension/flexion,
2. hip abduction/adduction,
3. hip external/internal rotation,
4. knee extension/flexion,
5. ankle plantarflexion/dorsiflexion,
6. ankle eversion/inversion.

For each DoF, there is a passive moment–angle relation. In addition, each DoF is consisted of two directions, whose movements are characterized by two active moment–angle relations and two active moment–velocity relations. Thus, we look to investigate a total of six passive relations, 12 isometric relations, and 12 isokinetic relations.

For each relation, we used a combination of keywords to initiate batch searches in Google Scholar with the software Publish or Perish. To guarantee search efficiency, we experimented multiple combinations of keywords in search of 13 target studies, which contain the desired moment data for ankle plantar-/dorsiflexion. The keyword set selected for the final batch search is one that led to the most target papers (12 out of 13) while having them appear in top search rankings (on average 20^th^ place in the list of 200 results), and it has the following format:

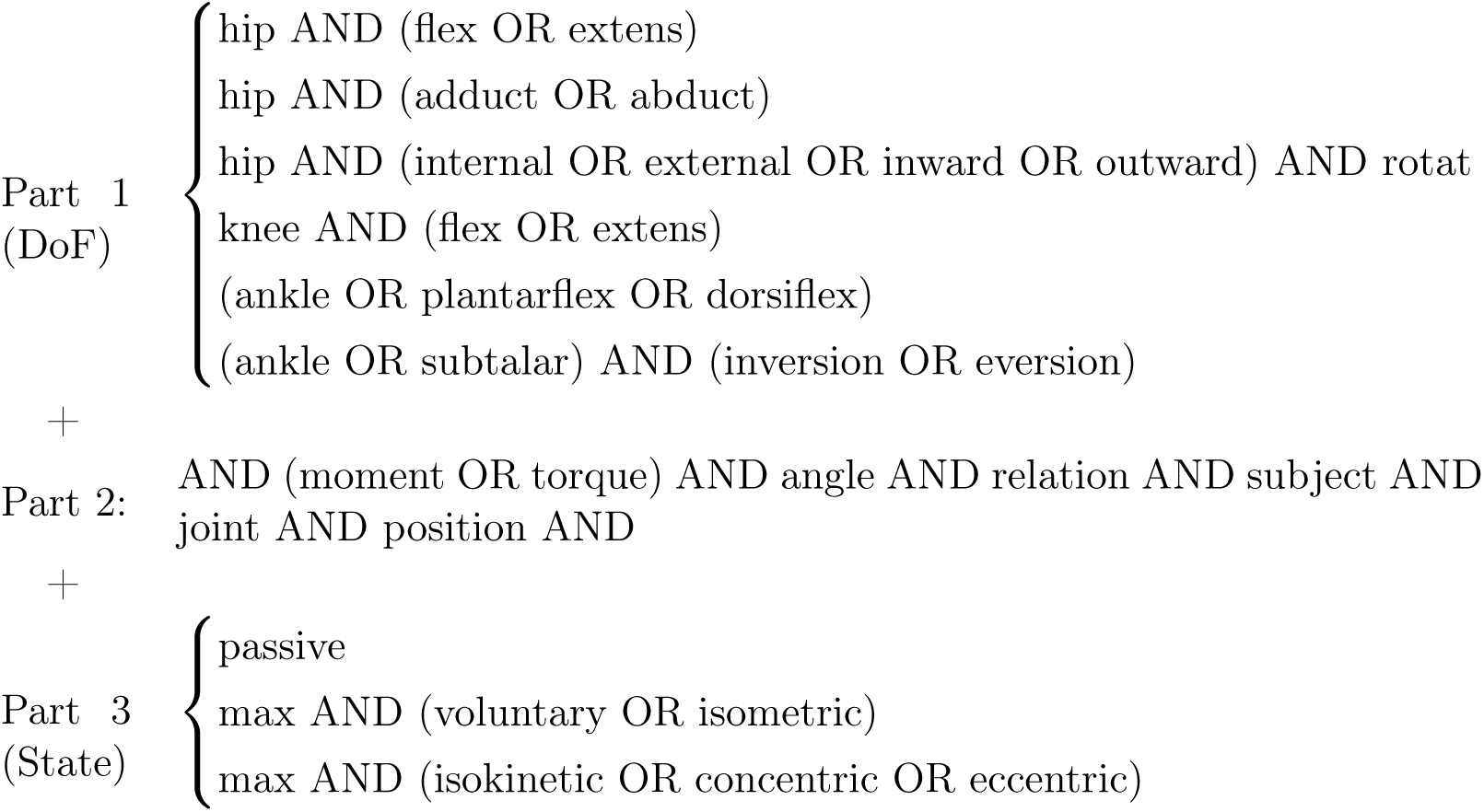

For all three sagittal DoFs and hip ab-/adduction, which are considered more dominant in motions, we examined the first 500 search results for the isometric relation and 200 each for the passive and isokinetic relations. For the other two DoFs, only the first 200 results for the isometric relation and 100 each for passive and isokinetic were examined. Other details of data collection are shown in Fig 1 and can also be found in Appendix A. Note that we also initiated searches oriented towards muscle moment arms for another study, but the search results were examined altogether. This serves as a compensation for the potential overfitting of the keywords, in case some records are missed in the searches intended for moment data.

**Figure 1.**
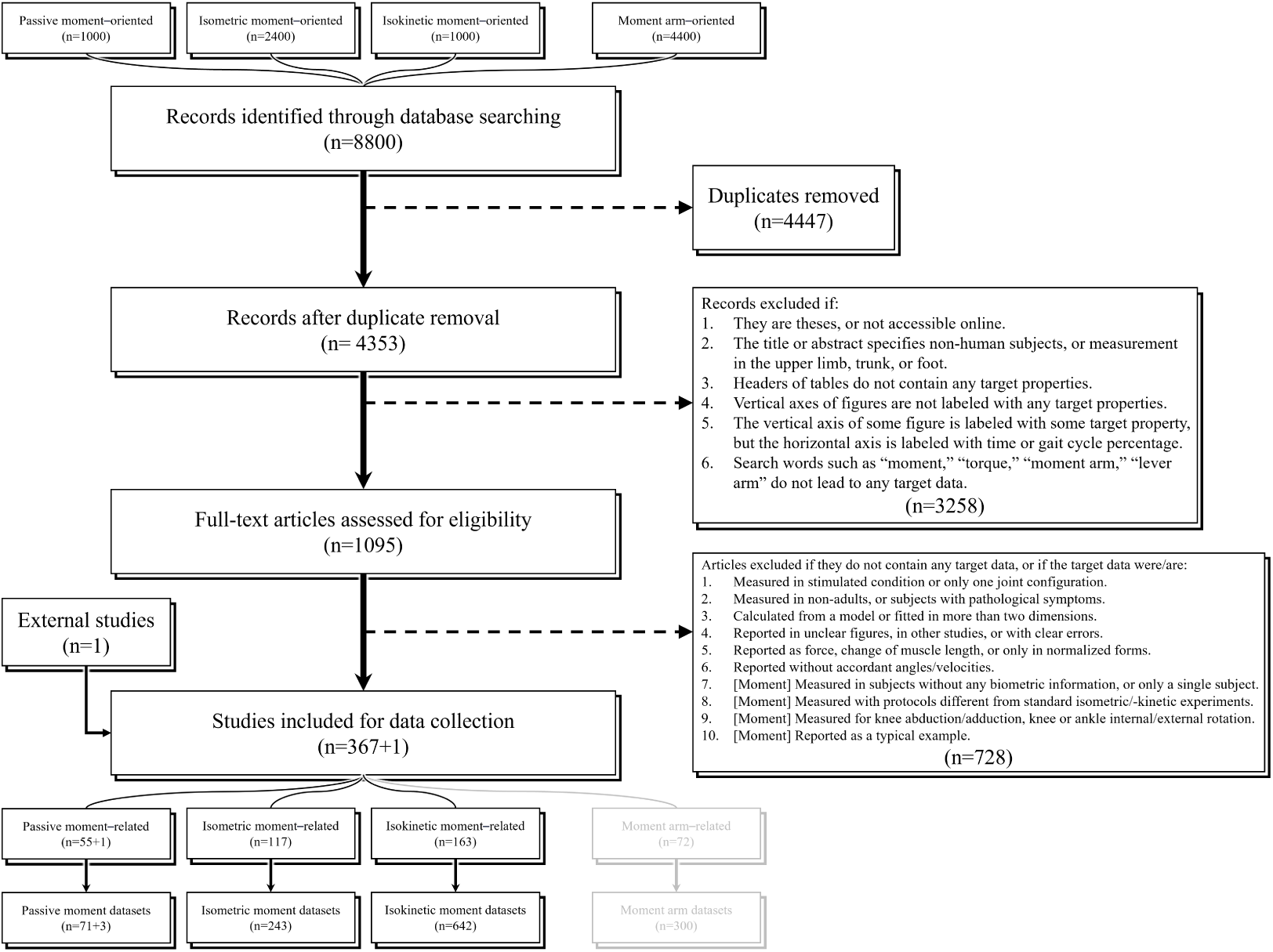
Workflow of study selection. Moment arm–related datasets are not presented in this study, but records identified by searches intended for moment arm data were examined for moment data.

In data collection, if a study presents its data in graphs rather than in numbers, we use Graph Grabber for digitization. The data are categorized by attributes listed in Table 1, and a dataset is defined as a set of measurements with distinction in either its reference, subject information, measured motion, or data type; in a few exceptional cases, datasets are distinguished by the tertiary DoF. In each dataset, there is at least one curve, which is comprised of measurements from at least two joint configurations, and some datasets contain multiple curves, each comprised of multiple measurements. The data for each curve are stored as a 3-column matrix, with the first, second, and third column for the angles in the primary and secondary DoFs, and the moments. The sign rule for angle and moment in each DoF is based on the ISB recommendations (Wu et al., 2002; Derrick et al., 2020) except for knee flexion/extension, which is reversed so that anti-gravity motions share the negative sign. The sign rule for velocity follows after Hill (1938), where concentric is defined as positive. If the angle in the secondary DoF is not specified in a study, we label it as NaN.

**Table 1.**
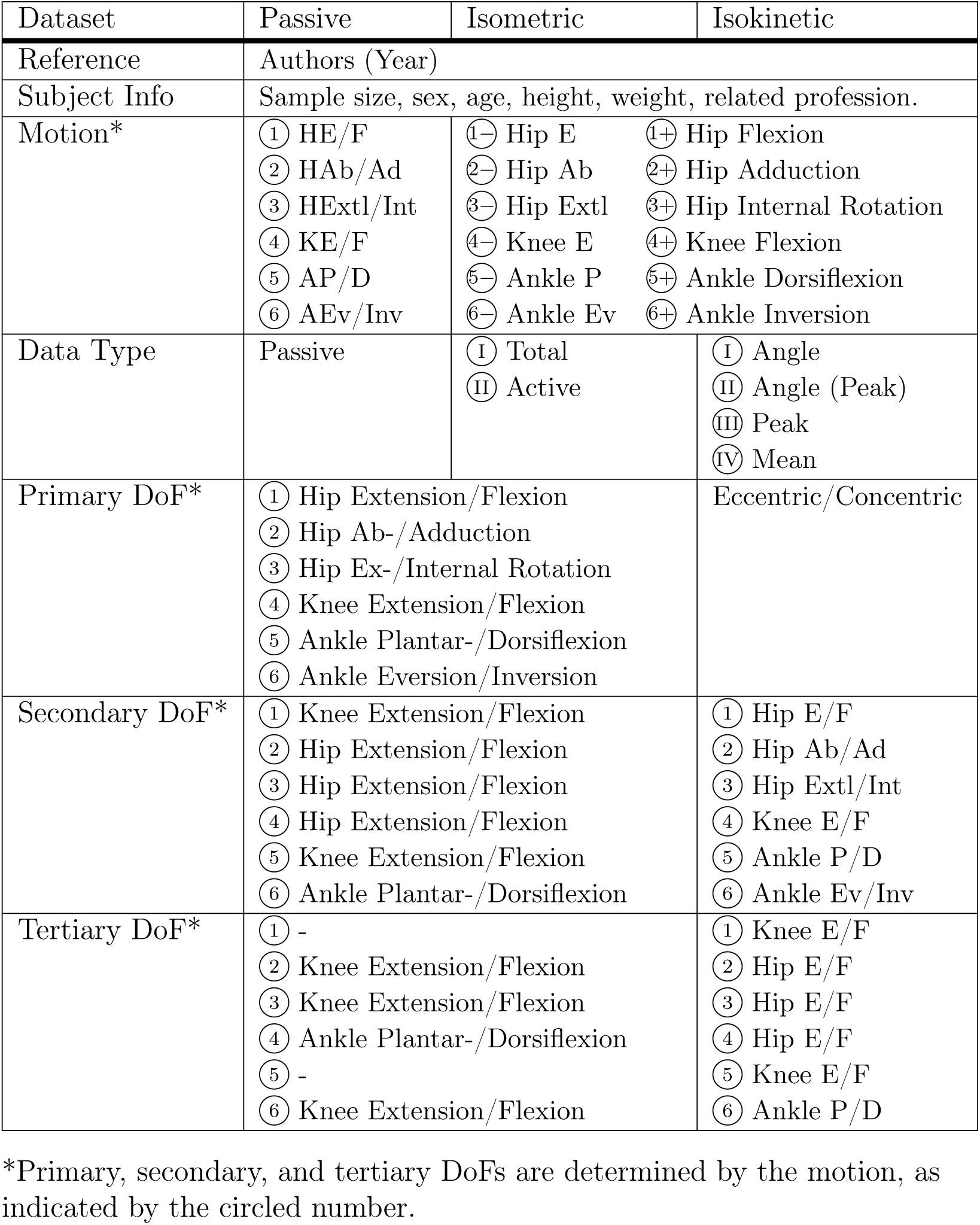
Metadata for passive, isometric, and isokinetic datasets.

Due to the massive size of the gathered datasets, we used 3D and 2D plots as a means of visualization to present the data. Importantly, since the data are multidimensional and the measured joint configurations are different across datasets, typical methods of meta-analysis no longer apply, and we did not proceed to combine the results. Also, considering the heterogeneity between studies, we refrain from making conclusive statements for any motion on the representative range and magnitude of joint angle or moment. For biomechanical analysis, readers are encouraged to select datasets with characteristics of subjects and means of measurement conforming to their object of study.

For the potential need of normalization and comparative analysis, we specified the sample size, sex, age, height, and weight of subjects, as well as the information of kinetics-based professions, such as solider, dancer, and athlete. We also marked if isometric data were corrected of the passive moments, and if isokinetic data were obtained as angle-specific, peak, or mean values within the test range of motion (RoM). For convenience, we provide a catalog with details of the datasets (Appendix A). In isokinetic datasets, if the moment data are peak or mean values, the angle in the secondary DoF (the moving DoF) is labeled as NaN, but the tested RoM is recorded in our catalog if specified in the original study.

## 3. Results

As shown in Fig 1, a total of 367 studies were identified from 4353 records searched in the database. Along with the one target paper which failed to show up in the pilot search, the studies yielded 74 passive datasets, 243 isometric datasets, and 642 isokinetic datasets. Table 2 shows the distribution of datasets for each motion. See Appendix B for a summary of collected studies and datasets.

**Table 2.**
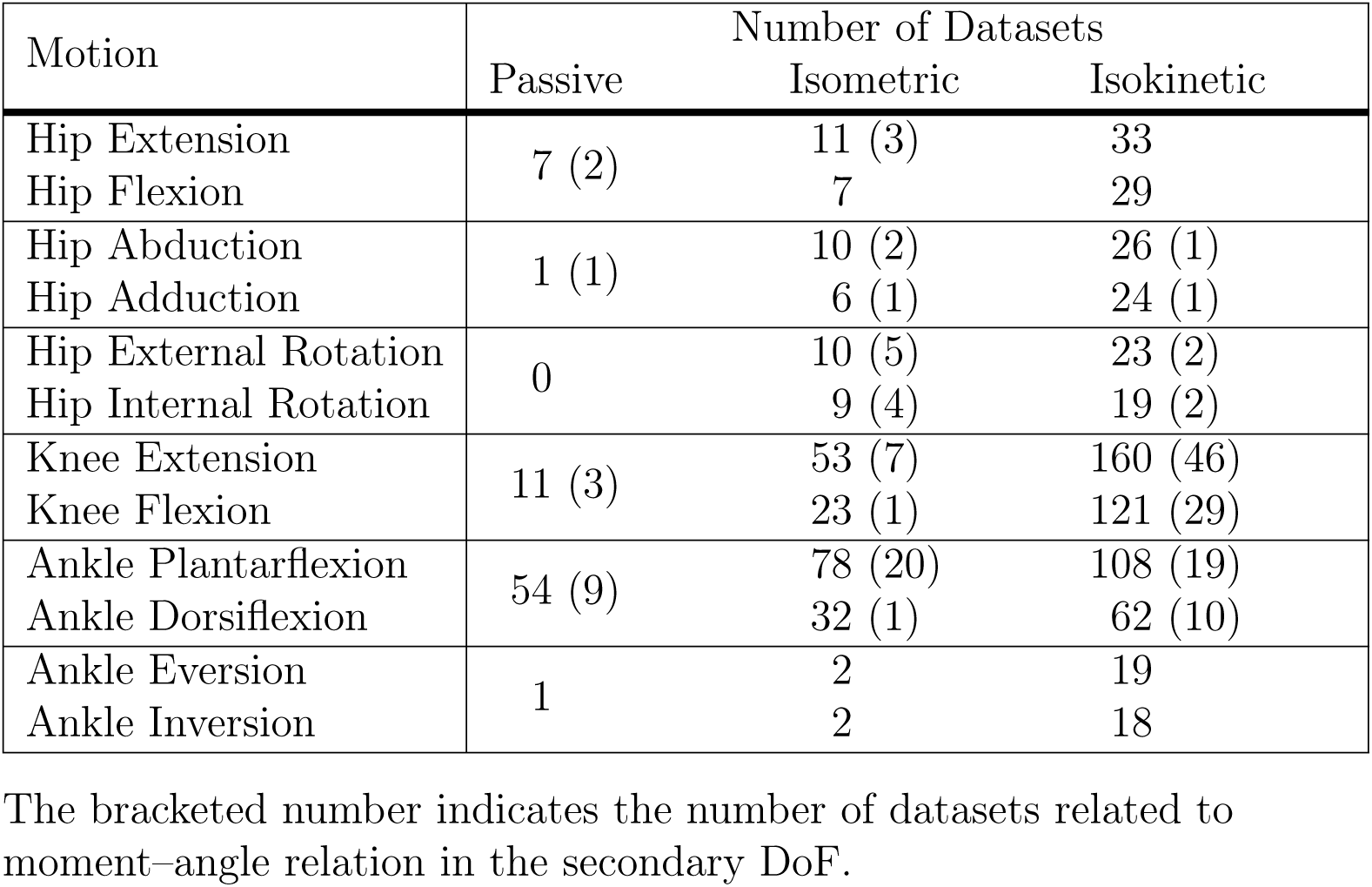
Distribution of datasets for each motion.

The gathered datasets are visualized in 3D and 2D plots (Figs 2–17). The age and sex of the subject group are indicated by the shape and color of the scatters: circle (20–39 yr), square (40–59 yr), triangle (*≥* 60 yr), red (predominantly female), blue (predominantly male), purple (mixed). If the angles for the secondary DoF are not specified in the original study, the dataset is plotted with dotted line(s) and unfilled scatters. Note that some datasets are not normalized and shown in 2D plots if subject height and weight are not specified in the original work.

## 4. Discussion

### 4.1. Study Heterogeneity

Before looking at the results in detail, we begin with the heterogeneity in the collected studies to explain for some of the variations observed in the moment datasets.

It is well-accepted that muscle strength is related to sex, age, and weight. Thus in the 2D plots, we normalized the moment data against the product of subject height and weight to decrease the range of variation (Handsfield et al., 2014). Although sex- and age-correlations are not a focus of this study, we labeled the curves with subject sex and age, as they might be associated with the variation in the normalized results. Also, we included datasets obtained from professions related to physical performance, but note that practicing such a profession could change the kinetic characteristics (Cometti et al., 2001; Moltubakk et al., 2018; Oliveira and Gonçalves, 2009; Ueno et al., 2018). For clarity of the figures, we did not include this information in the plots but it is recorded in the metadata.

Some other parts of the variation may come from the experiment apparatus and protocol. Before the prevalence of commercial dynamometers such as Cybex, Biodex, KinCom, and Isomed2000, researchers customized devices for moment measurement (Marsh et al., 1981; Smidt, 1973; Wiemann and Hahn, 1997), and this variety of apparatus makes it complicated to compare between studies. For instance, perhaps for the consideration of convenience, researchers would use hand-held dynamometers to measure force and calculate moment with manually-measured lever arm, but this method would yield different results compared with the conventional isokinetic dynamometer (Bazett-Jones and Squier, 2020) or if different techniques are used (Krause et al., 2014). There are more variables in terms of the protocol: e.g., lying in supine or prone (Barr and Duncan, 1988; Mańka et al., 2022; Turpin et al., 2014), measurement by side or leg dominance (T’Jonck et al., 1997; Maganaris et al., 1998), provision of visual feedback (Hald and Bottjen, 1987), and various ways of stabilization (Alt et al., 2016; Lavin and Gross, 1990; Magnusson et al., 1993; Nisell et al., 1989; Weir et al., 1996a). Even clenching of the teeth or positioning of the tongue are reported to induce significant differences in measurements (di Vico et al., 2013; Sasaki et al., 1998). Importantly, the center of rotation is critical as it directly affects measurement accuracy, but how this center is determined is not typically described, and one can expect miscellaneous approaches from studies.

For most part, the datasets we gathered inevitably suffer from the impact of the factors mentioned above as well as many not mentioned, and the curves plotted in Figs 2–11, 13–16 are not particularly consistent in terms of the magnitude. Our meta-visualization serves as a reminder of the potential errors in individual studies and offers a more general reference for those who wish to perform biomechanical analysis with moment data. As will be shown in later discussion, it is easy to end up in distinct conclusions when only a few datasets are used in the representation of joint kinetic characteristics.

### 4.2. Passive Moment

Passive moment is an elementary kinetic property in the joint: Its exertion is mostly mechanical, hence easy to measure, and it is also part of the *total* moment measured with activated muscles, so it is often corrected for when studying active mechanics.

In the lower limb, most of the datasets we gathered are for the sagittal plane, especially ankle plantar-/dorsiflexion (Table 2). Lower limb motion in the sagittal plane has been a major interest in the biomechanics community, for their dynamics are more dominant in daily locomotion such as walking and running, and the dynamometers used to measure joint moment are generally designed to have the subjects stabilized in the sagittal plane. So it is natural that researchers tend to report sagittal moment data. Especially considering that the RoM and muscle moment arm are generally larger in the sagittal plane (*citation*: companion submission), the stretch of passive components will be more intensive, thus it is more necessary and convenient to measure the passive moment.

The pattern of passive moment–angle relation is simple, because it mainly depends on the mechanical stretch of muscles in different joint positions. In theory, there should be an increase of extension moment as the joint flexes and an increase of flexion moment as the joint extends. Meanwhile, due to the existence of biarticular muscles such as the rectus femoris, hamstrings, and gasctronemii, this rule is reversed for the adjacent joint: there should be an decrease of extension moment as the adjacent joint extends and an decrease of flexion moment as the adjacent joint flexes.

We start from the passive moment–angle relations in the secondary DoF (Figs 2–4, left), that is how the passive moments in the hip, knee, and ankle change according to joint angles at the knee, hip, and knee. In the hip, although only two studies contain relevant data, we can still see a decrease of hip extension moment or an increase of hip flexion moment as the knee flexes, especially evident when hip is highly extended or flexed (Fig 2, left). A similar pattern is observed in the knee, but with a smaller magnitude (Fig 3, left); this could be because the difference of biarticular muscles in their moment arms about the hip and knee (*citation*: companion submission). In the ankle, the decrease of plantarflexion moment along the increase of knee flexion is evident when the ankle is dorsiflexed (Fig 4, left), and the magnitude is quite large, consistent with the fact that the gastrocnemii constitute one third of the triceps surae (Handsfield et al., 2014; Ward et al., 2009).

**Figure 2.**
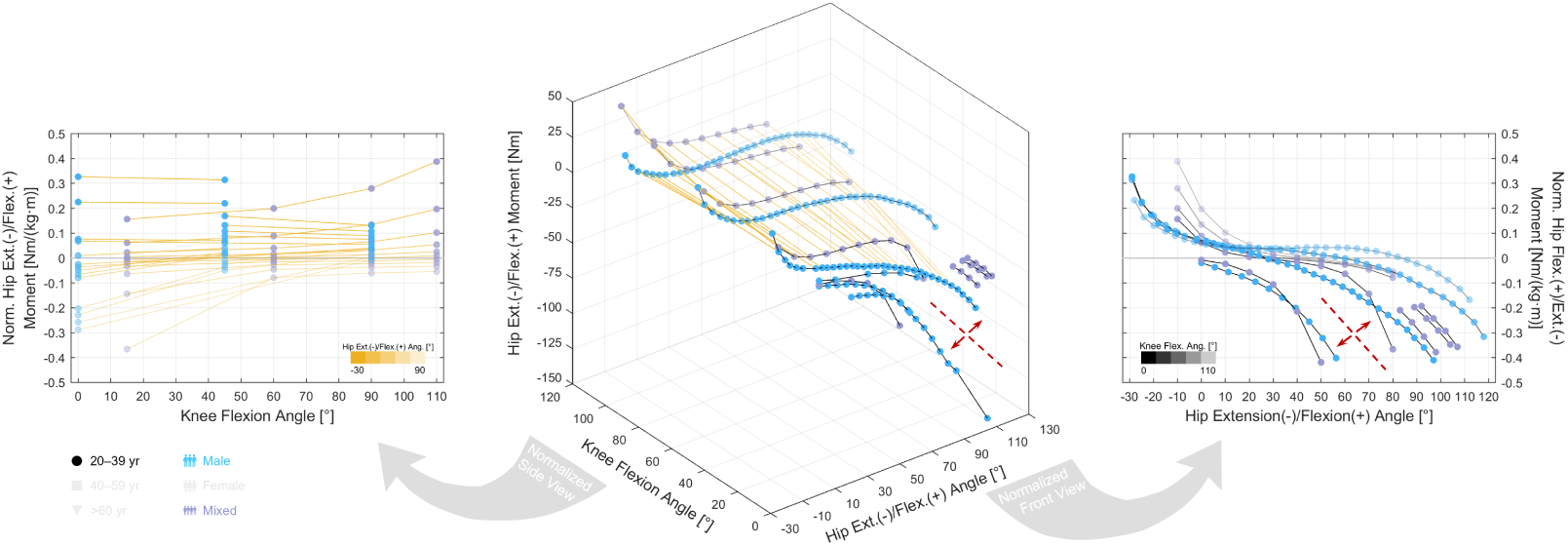
Relations of passive hip extension/flexion moments. Left: Normalized 2D relation with knee flexion angle. Center: 3D relation with hip and knee flexion angles. Right: Normalized 2D relation with hip flexion angle. Data are normalized in the 2D plots by the product of subject height and weight. The transparency of the curve segment denotes the angle in the relatively fixed DoF (color bar): with yellow for hip flexion and black for knee flexion. Notice the curves may be divided into two groups as indicated by the red dashed line and arrows.

**Figure 3.**
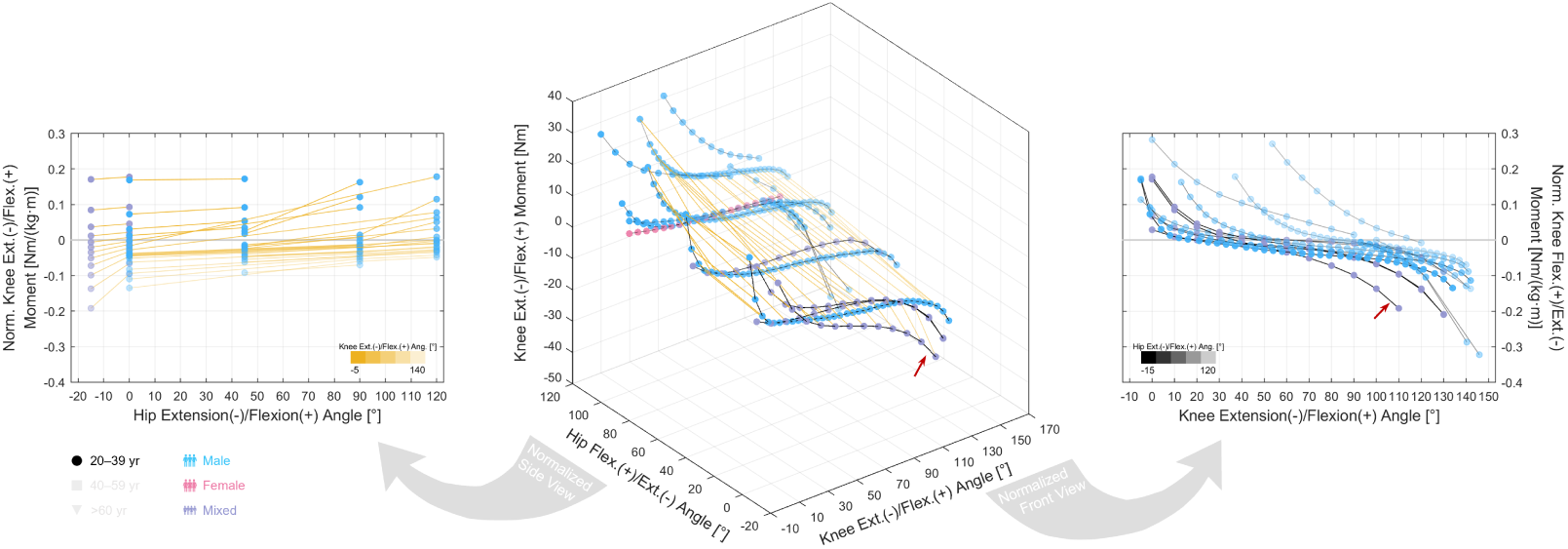
Relations of passive knee extension/flexion moments. Left: Normalized 2D relation with hip flexion angle. Center: 3D relation with knee and hip flexion angles. Right: Normalized 2D relation with knee flexion angle. Data are normalized in the 2D plots by the product of subject height and weight. The transparency of the curve segment denotes the angle in the relatively fixed DoF (color bar): with yellow for knee flexion and black for hip flexion. Notice the curve pointed by the red arrow is not plotted in the plane of 0° hip flexion, hence differs from other curves with little color transparency.

**Figure 4.**
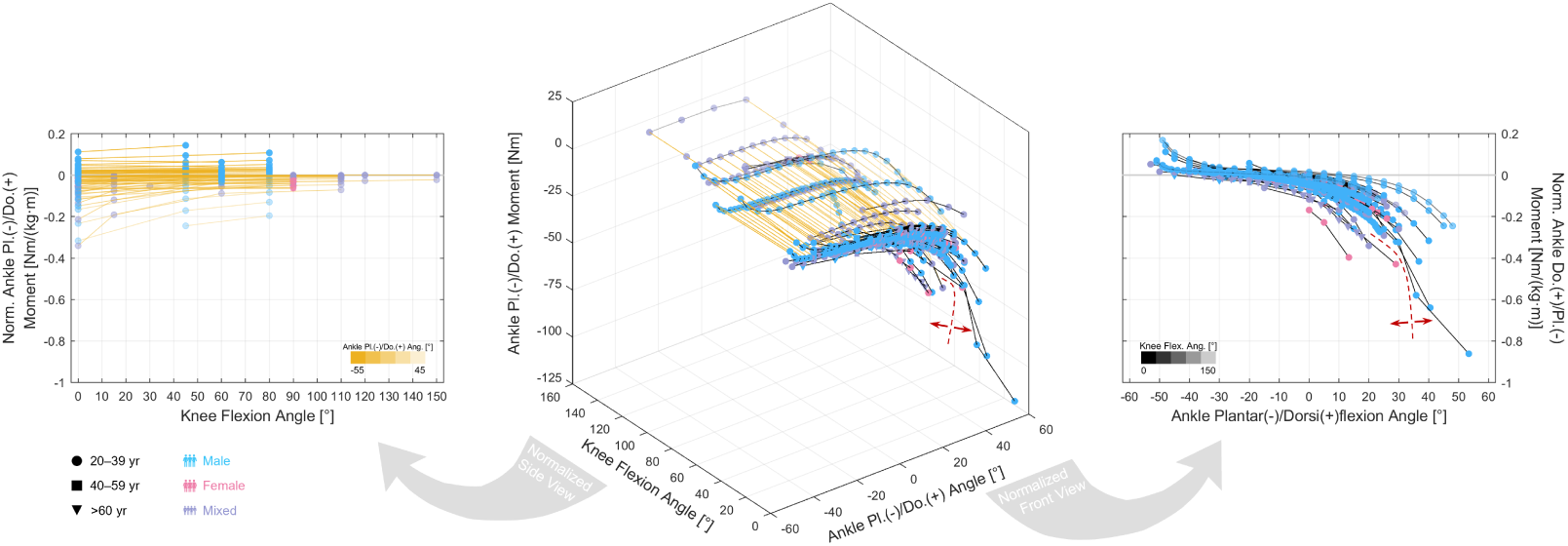
Relations of passive ankle plantar-/dorsiflexion moments. Left: Normalized 2D relation with knee flexion angle. Center: 3D relation with ankle dorsiflexion and knee flexion angles. Right: Normalized 2D relation with ankle dorsiflexion angle. Data are normalized in the 2D plots by the product of subject height and weight. The transparency of the curve segment denotes the angle in the relatively fixed DoF (color bar): with yellow for ankle dorsiflexion and black for knee flexion. Notice the curves may be divided into two groups as indicated by the red dashed line and arrows.

In the primary DoF, the pattern in the passive moment–angle curves is, as expected, generally exponential (Figs 2–4, center and right), but this simple pattern causes an issue in the precise depiction of passive moment– angle relations. Since passive moment increases much more in the latter half of RoM, the error in the horizontal axis will be magnified in the vertical axis. For example in Fig 2 (right), looking at the hip moment–angle curves with non-transparent colors (extended knee), it is easy to distinguish two groups: the *left* group starts from no moment at the neutral hip position to a maximum of 0.4 Nm/(kg*·*m) at 50*^◦^* hip flexion, while the *right* group has a zero-moment angle of 20*^◦^*–30*^◦^* and reaches a similar maximum only at 100*^◦^* hip flexion. This disparity is well worth noticing, because it is not just a shift in the zero-moment angle but really about how much passive moment is exerted by the hip in high flexion positions.

In Fig 2 (middle), one curve from the left group (Wiemann and Hahn, 1997) reaches 142 Nm at 105*^◦^* hip flexion, which is about three times the maximum of the curves in the right group. This is a difference of an entire order of magnitude, and if the left group is correct, then active hip moment measured in high flexion positions must be corrected of passive moment, otherwise the isometric and isokinetic hip moment–angle relations would be much distorted.

One reason behind such discrepancy could be the way researchers positioned the subjects in the so-called “neutral position,” which is usually supine. Wiemann and Hahn (1997) placed a “supporting jack” under the subject’s sacrum, which could have anteriorly tilted the pelvis when the thigh appeared to be in line with the trunk. If so, the measurement may have in fact started from a slightly flexed hip angle rather than zero, inducing a left shift of the curve. Miyamoto et al. (2017) had the subjects lying on their side, where the pelvis might no longer be posteriorly tilted by the weight of the spine and the back muscles, and the curve would shift similarly to the previous example. Halbertsma et al. (1999) subtracted from all data the measurement in the neutral position as the moment induced by limb and apparatus weight, assuming no passive moment in the neutral position. However, some studies reported flexion moment up to 10 Nm (Riener and Edrich, 1999; Silder et al., 2007), hence the curve might be shifted downwards, visually inducing a left shift.

A similar issue is observed in passive ankle plantar-/dorsiflexion moment curves (Fig 4, center and right). Unlike the hip or knee, there lacks a natural neutral position for the ankle, and the ankle angle is usually define as the included angle between the tibia and the sole of the foot. However, the tibia has thickness and is not a line, so there will be significant differences depending on whether the length axis is chosen as the anterior border (Toft et al., 1989), the posterior border (Marsh et al., 1981), the connection between the fibular head and medial malleolus (Weiss et al., 1986), or by some other standards. Furthermore, the RoM of ankle plantar-/dorsiflexion is much smaller than that of the hip and knee, so even a small variation of 10*^◦^* would appear huge for the passive moment-angle curve. Similar to hip extension/flexion, in Fig 4 (right), there is a *right* group of passive plantar-/dorsiflexion moment–angle curves (Savage et al., 2015), with no passive moment in the neutral position, and a *left* group, reporting a zero-moment angle of 10*^◦^*–15*^◦^* plantarflexion (Toft et al., 1989; Winegard et al., 1997).

In comparison, the knee is free of this issue, because full extension serves as the neutral position and the extent of hyperextension is small compared with the large RoM of knee extension/flexion. In Fig 3 (middle and right), the seemingly unmatched curves are mainly the results of different hip angles, and the curves in the 3D plot are magnitude-wise consistent in their respective *xz* -planes.

### 4.3. Isometric Moment

It is well-known from the muscle force–length relation that maximal muscle force changes varies with muscle length, and there is an optimal length where maximal muscle force reaches its peak. Hence it is reasonable to expect a similar shape for the joint moment–angle relation, where there is also a peak value and an accordant optimal joint angle.

Nevertheless, based on our gathered datasets, although maximal joint moment does operate on the ascending limb, a plateau is not typically reached and it can be hard to identify a peak value. Overall, the moment–angle curves in the primary DoF exhibit a similar pattern: When the data for two directions of a DoF are plotted together, the two groups of curves are visually similar to an equal sign rotated clockwise (Figs 5–9, top right). In other words, the maximal moment exerted in either direction of a DoF increases as joint angle increases along the other direction, where the agonists are lengthened; we refer to it as *parallel ascend*.

**Figure 5.**
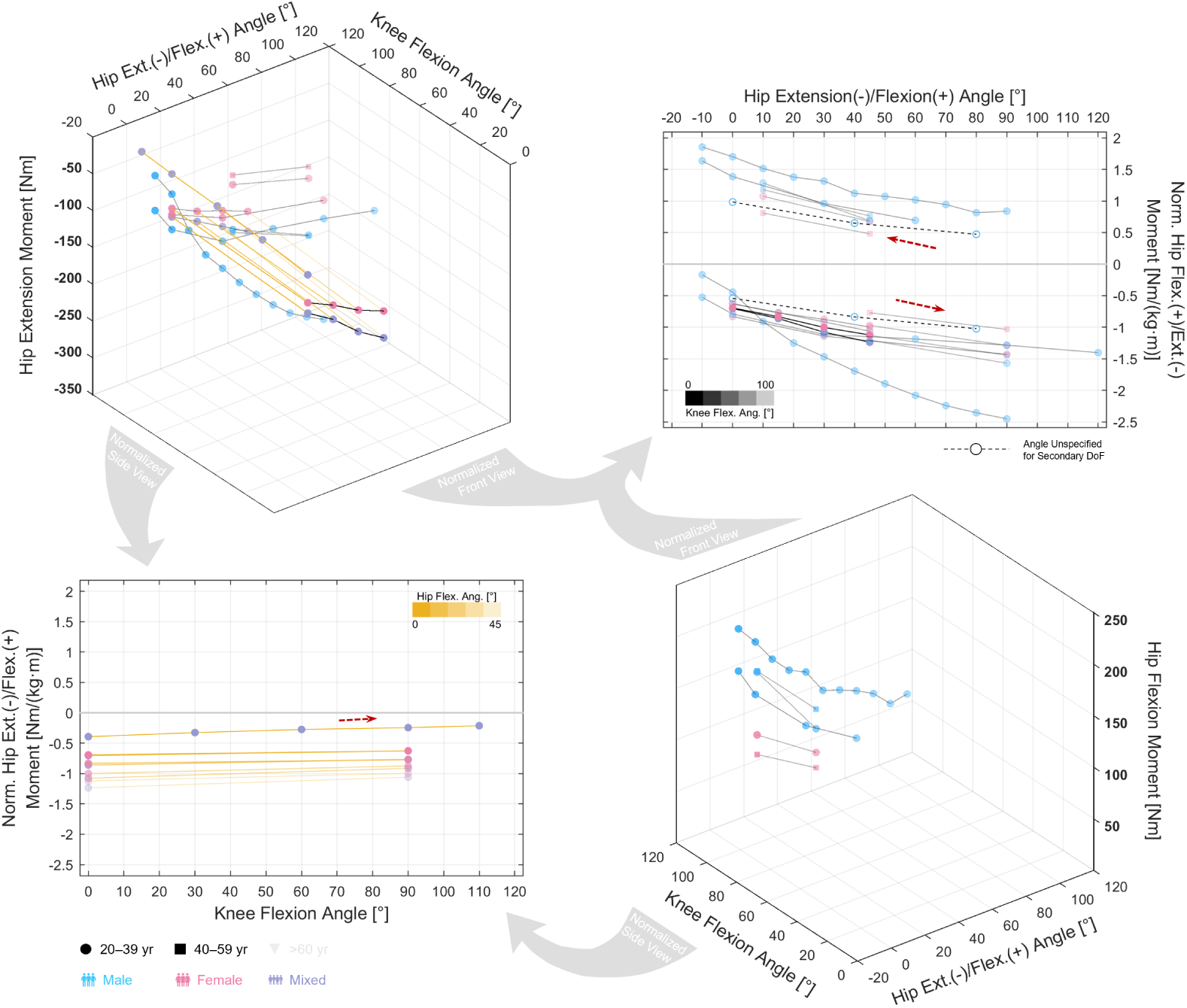
Relations of isometric hip extension/flexion moments. Top left: 3D relation of extension moment with hip and knee flexion angles. Top right: Normalized 2D relation with hip flexion angle. Bottom left: Normalized 2D relation with knee flexion angle. Bottom right: 3D relation of flexion moment with hip and knee flexion angles. Data are normalized in the 2D plots by the product of subject height and weight. The transparency of the curve segment denotes the angle in the relatively fixed DoF (color bar): with yellow for hip flexion and black for knee flexion. The direction of muscle stretch is indicated by the red arrows.

**Figure 6.**
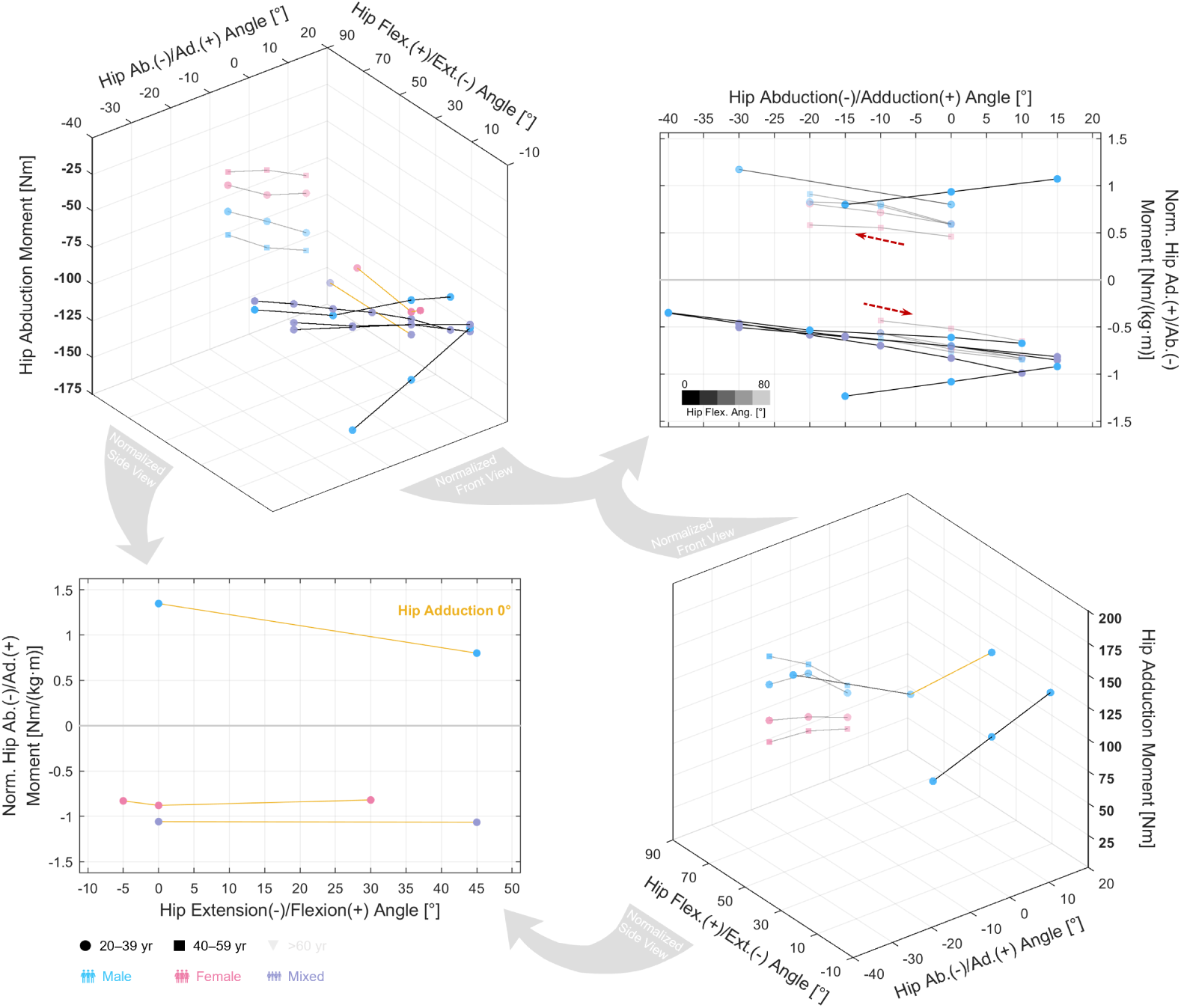
Relations of isometric hip ab-/adduction moments. Top left: 3D relation of abduction moment with hip adduction and flexion angles. Top right: Normalized 2D relation with hip adduction angle. Bottom left: Normalized 2D relation with hip flexion angle. Bottom right: 3D relation of adduction moment with hip adduction and flexion angles. Data are normalized in the 2D plots by the product of subject height and weight. The transparency of the black curve segment denotes hip flexion angle (color bar). The direction of muscle stretch is indicated by the red arrows.

**Figure 7.**
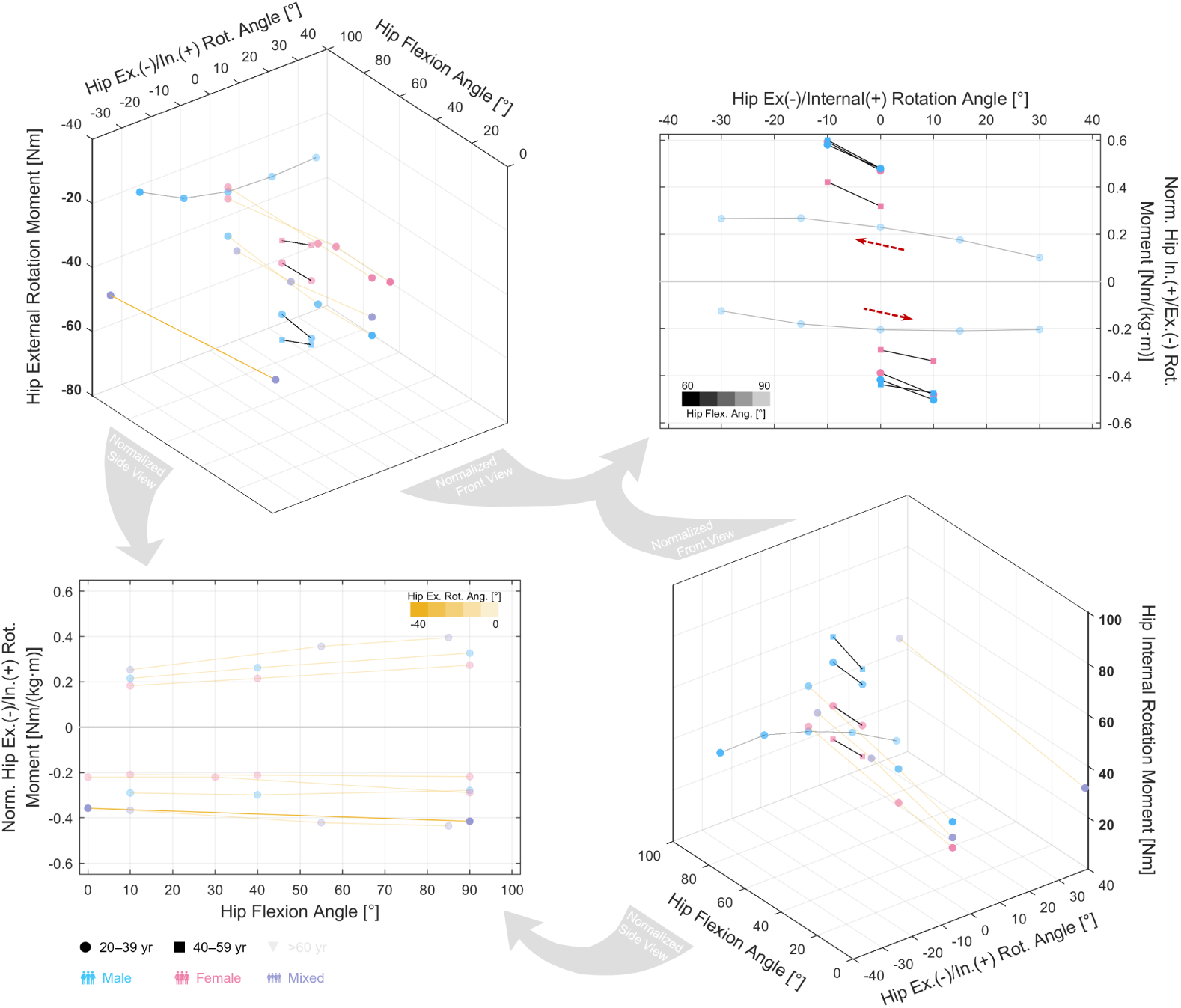
Relations of isometric hip ex-/internal moments. Top left: 3D relation of external rotation moment with hip internal rotation and flexion angles. Top right: Normalized 2D relation with hip internal rotation angle. Bottom left: Normalized 2D relation with hip flexion angle. Bottom right: 3D relation of internal rotation moment with hip internal rotation and flexion angles. Data are normalized in the 2D plots by the product of subject height and weight. The transparency of the curve segment denotes the angle in the relatively fixed DoF (color bar): with yellow for hip external rotation and black for hip flexion. The direction of muscle stretch is indicated by the red arrows.

**Figure 8.**
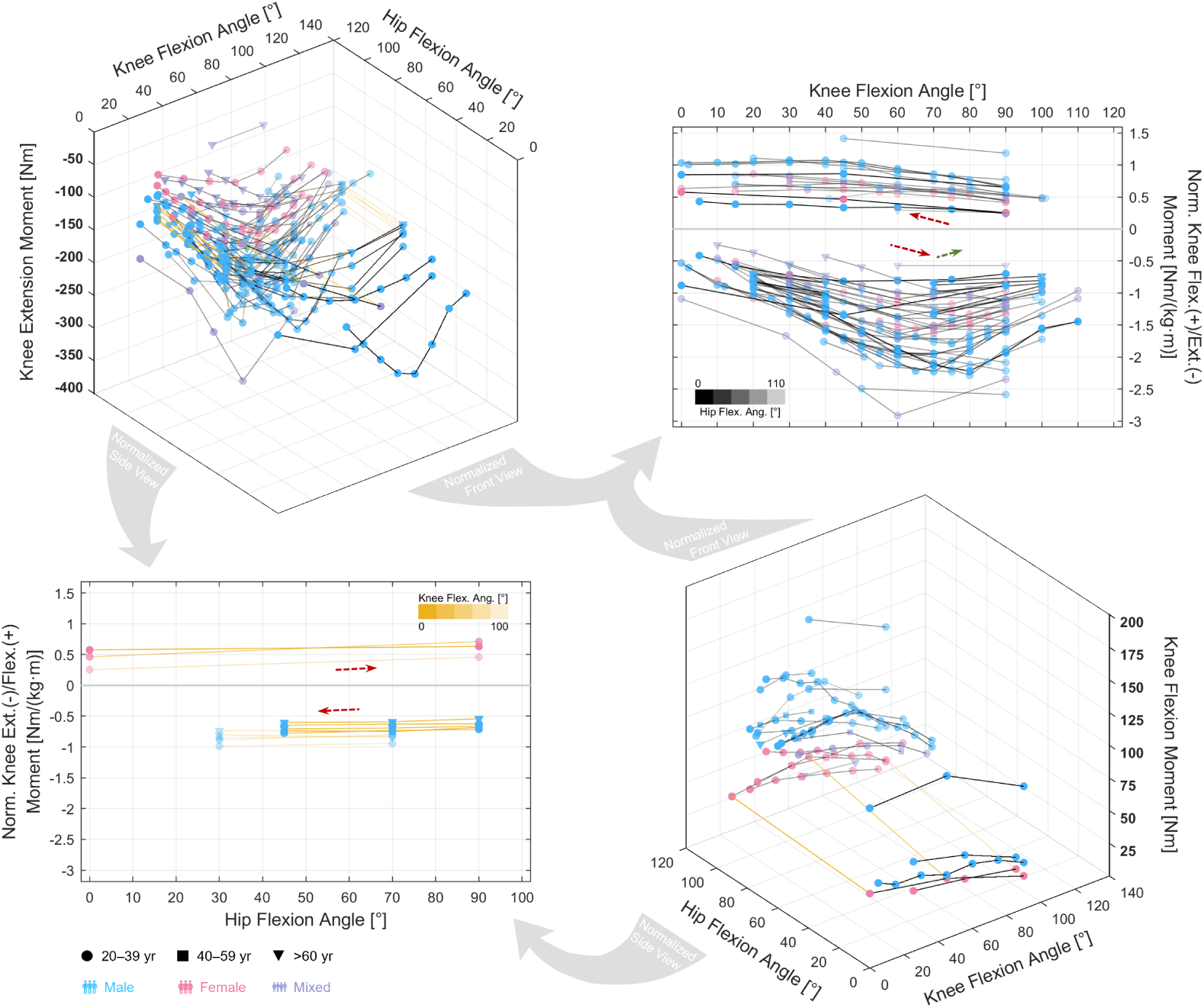
Relations of isometric knee extension/flexion moments. Top left: 3D relation of extension moment with knee and hip flexion angles. Top right: Normalized 2D relation with knee flexion angle. Bottom left: Normalized 2D relation with hip flexion angle. Bottom right: 3D relation of flexion moment with knee and hip flexion angles. Data are normalized in the 2D plots by the product of subject height and weight. The transparency of the curve segment denotes the angle in the relatively fixed DoF (color bar): with yellow for knee flexion and black for hip flexion. The direction of muscle stretch is indicated by the red arrows.

**Figure 9.**
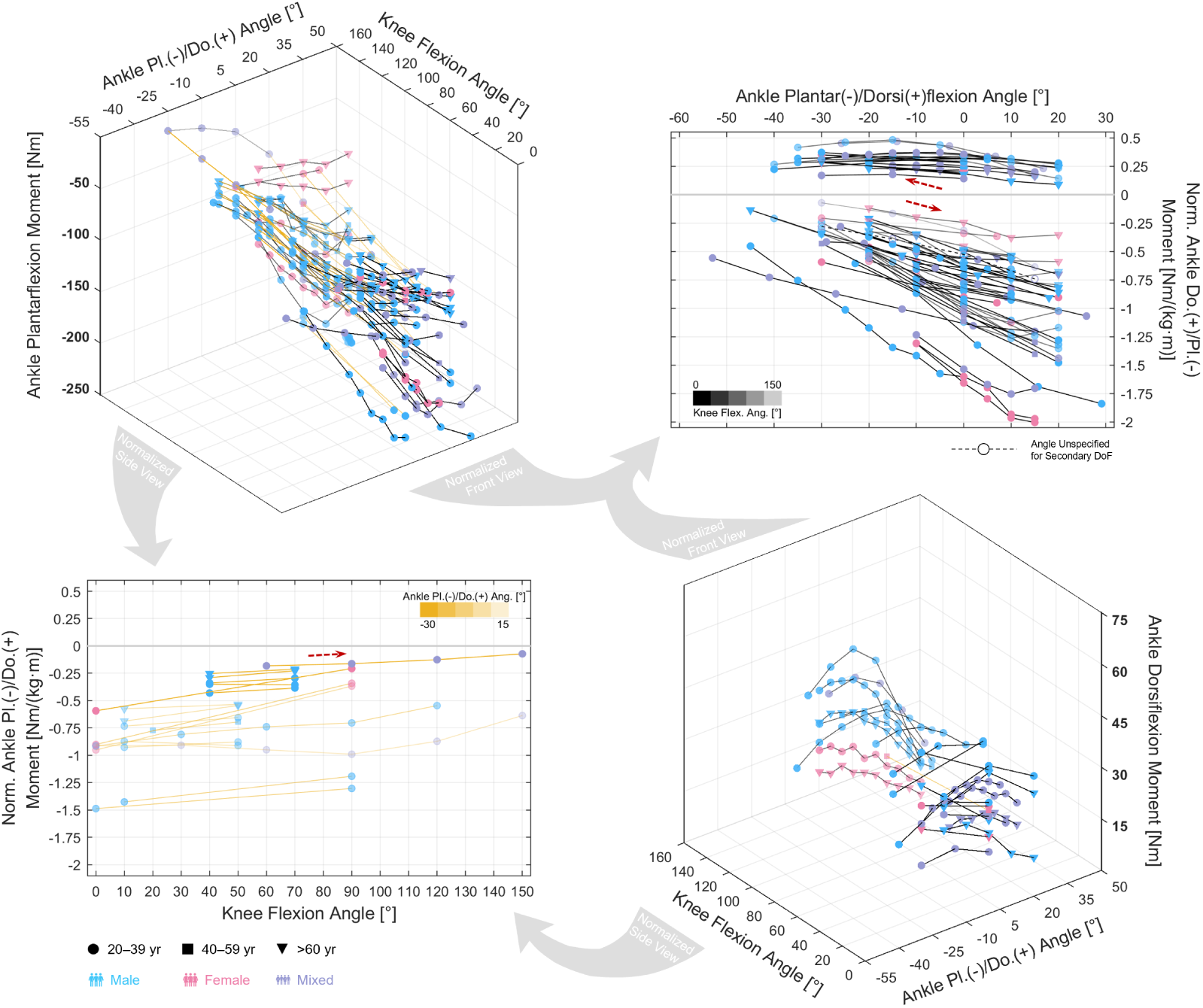
Relations of isometric ankle plantar-/dorsiflexion moments. Top left: 3D relation of plantarflexion moment with ankle dorsiflexion and knee flexion angles. Top right: Normalized 2D relation with ankle dorsiflexion angle. Bottom left: Normalized 2D relation with knee flexion angle. Bottom right: 3D relation of dorsiflexion moment with ankle dorsiflexion and knee flexion angles. Data are normalized in the 2D plots by the product of subject height and weight. The transparency of the curve segment denotes the angle in the relatively fixed DoF (color bar): with yellow for ankle dorsiflexion and black for knee flexion. The direction of muscle stretch is indicated by the red arrows.

This is not at all a new finding but rather easily expected. When a joint approaches full extension, there should be less need to further extend the joint, because there simply is no more space for extension, and it should be more likely that flexion is to be initiated; otherwise human kinematics would easily get stuck in some extreme position. Thus from the perspective of energetic efficiency, higher flexion moments and lower extension moments are needed in extended positions and vice versa. It seems this is indeed where evolution brings us to.

For hip extension/flexion, when normalized against subject height and weight, the magnitude of moment is rather consistent between studies (Fig 5, top right): Hip flexion moment is 1.5–2.0 Nm/(kg*·*m) in extended and slightly flexed positions, and decreases below 1.0 Nm/(kg*·*m) as the hip flexes. Hip extension moment is 1.0–1.6 Nm/(kg*·*m) in highly flexed positions, and decreases below 1.0 Nm/(kg*·*m) as the hip extends; Anderson and Pandy (1999) reported extension moment exceeding 2.0 Nm/(kg*·*m) in flexed positions, but the pattern is the same. In a similar fashion, hip adduction moment increases as the hip abducts, while abduction moment increases as the hip adducts (Fig 6, top right); with the exception of Anderson and Pandy (1999). Hip internal moment increases as the hip externally rotates, while external moment increases as the hip internally rotates (Fig 7, top right).

For knee flexion and ankle dorsiflexion, the ascent is less obvious (Figs 8 and 9, top right) because their magnitudes are much smaller than those of their counterparts, but it can be well observed when plotted alone, as shown in Figs 8 and 9 (bottom right). For knee extension, a peak at 60*^◦^*–80*^◦^* is observed, beyond which extension moment decreases, and the magnitude of this peak is 1.0–2.0 Nm/(kg*·*m). For ankle plantarflexion (Fig 8, top right), though the ascent is clear, the maximal moment in high dorsiflexion positions is 0.5–1.5 Nm/(kg*·*m), and several curves even reach 2.0 Nm/(kg*·*m). This variance is much larger than measurements in knee extension, and the conclusion stands even when passive moment are accounted for (Fig 10). One reason could be the aforementioned issue of the neutral position: Since there is a large error in determining the zero ankle angle, the moment–angle curve may be shifted horizontally.

**Figure 10.**
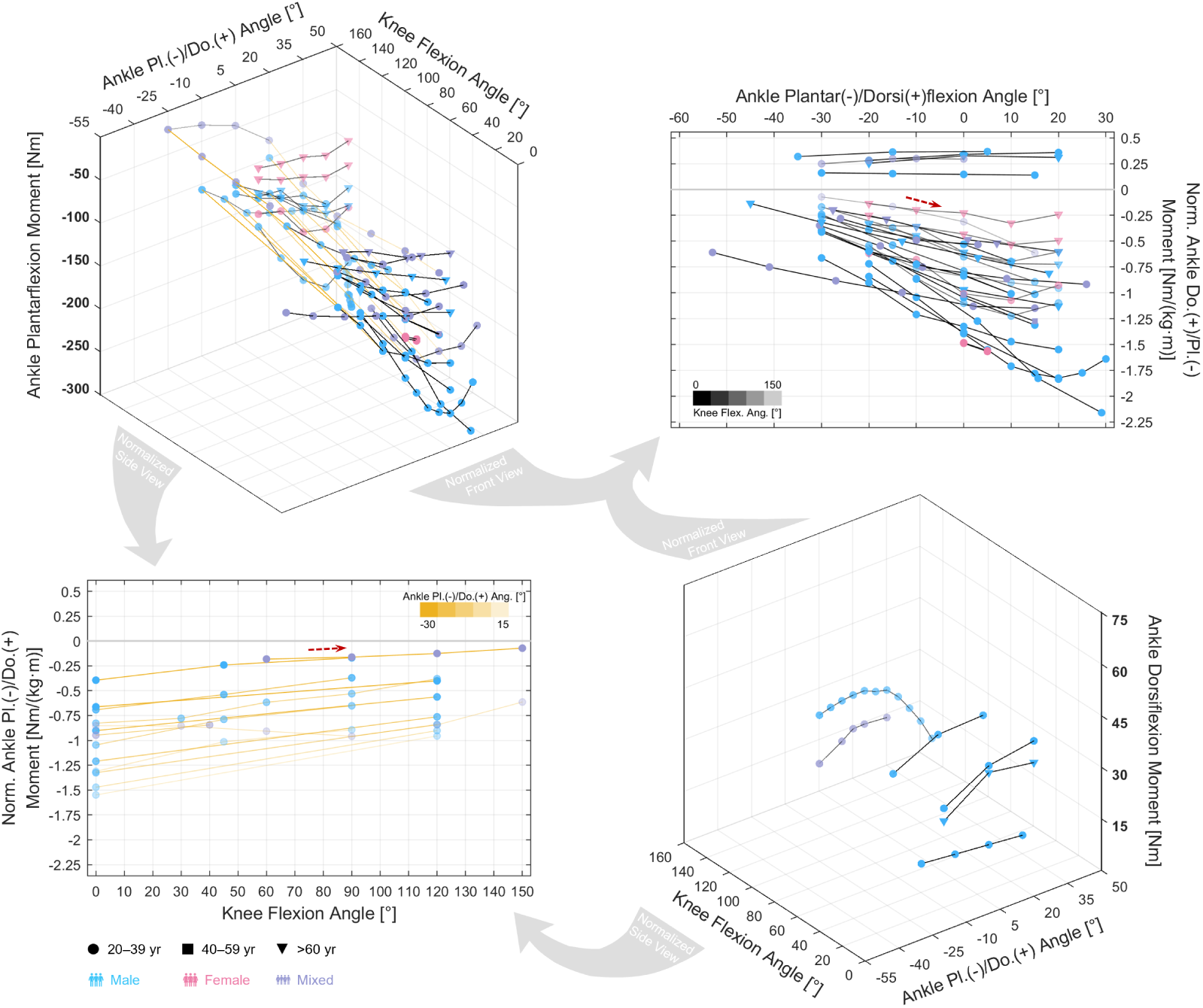
Relations of isometric ankle plantar-/dorsiflexion moments (passive corrected). Top left: 3D relation of plantarflexion moment with ankle dorsiflexion and knee flexion angles. Top right: Normalized 2D relation with ankle dorsiflexion angle. Bottom left: Normalized 2D relation with knee flexion angle. Bottom right: 3D relation of dorsiflexion moment with ankle dorsiflexion and knee flexion angles. Data are normalized in the 2D plots by the product of subject height and weight. The transparency of the curve segment denotes the angle in the relatively fixed DoF (color bar): with yellow for ankle dorsiflexion and black for knee flexion. The direction of muscle stretch is indicated by the red arrows.

As explained, measurements of joint moment may be influenced by a vast range of factors, and for this reason we try not to judge if a dataset is correct. However, it is still possible, and necessary, to evaluate if some are less reliable. Unlike density of blood cells or length of a sarcomere, maximal moment of a joint is not entirely a physical property whose magnitude requires direct measurements to determine. Ultimately, joint moments are for generating kinematics, and the moment required to generate simple locomotion is a bottom line for maximal moment output. For example, based on the kinematics of walking at normal speed (Camargo et al., 2021), it is has been calculated that a plantarflexion moment of 1.0–1.5 Nm/kg, or roughly 0.6–0.9 Nm/(kg*·*m), is required in the push-off phase, where the ankle is dorsiflexed. Therefore it is fair to question a dataset if the maximal plantarflexion moment measured in dorsiflexed positions is within or below this range, especially if healthy young subjects were tested.

For this reason, we specified in Fig 1 the exclusion of data fitted in more than two dimensions. Fitting a group of data into a 3D surface would inevitably smooth its peaks, and the results would risk contradicting the kinematics. For example, when fitting moment data to a joint angle–velocity surface, Anderson et al. (2007) estimated a maximal isometric moment of 95 Nm for their young subject group, roughly equivalent to 0.7 Nm/(kg*·*m). This value is barely enough for walking and completely insufficient for running (1.2 Nm/(kg*·*m); Fukuchi et al., 2017). If this dataset is used to calibrate a musculoskeletal model, even the simulation of locomotion may not be accurate. Hence, despite the labor-intensive experiments and valuable discoveries, the work of Anderson et al. (2007) and others (Khalaf et al., 2000; Hussain and Frey-Law, 2016; Marshall et al., 1990) are not included in this study.

Although our proposed method of comparing estimates of maximal joint moments to those required for locomotion is simple and straight-forward, we do not recommend using more intensive movements, such as jumping and squatting, as a more strict criterion for filtering. In isometric experiments, although subjects are instructed to contract their agonists as much as possible, the activation of agonists is not necessarily maximal (Mohamed et al., 2002; Onishi et al., 2002; Salzman et al., 1993; Worrell et al., 2001), nor is the activation of antagonists necessarily zero (Aagaard et al., 2000; Billot et al., 2011; Kellis and Baltzopoulos, 1997; Maganaris et al., 1998). It could be that the measurements obtained in a lab are always lower than the true maximal isometric moment, and too high a threshold could potentially eliminate too many experiments properly conducted.

Regarding the relations of moments in the sagittal plane (Figs 5, 8, 9, bottom left) in the secondary DoF, the pattern is similar to that exhibited in passive moment relations (Figs 2–4, left) and appears to be in *parallel descent* when plotted together (Fig 8, bottom left). For example, maximal isometric plantarflexion moment decreases as the knee flexes (Fig 9, bottom left), since the shortening of the gastrocnemii not only decreases its passive force but also brings active force down in the ascending limb of force–length curve. For moments in the non-sagittal planes, there are not enough data to draw any conclusions, but we do not expect to see parallel descent. This is because, unlike in the sagittal plane where biarticular muscles have opposite functions in the adjacent joints, the functions of a muscle in the sagittal and non-sagittal planes are not bonded. For instance, while both parts of the gluteus maximus function as hip extensors, the upper part is a hip abductor while the lower part is a hip adductor (Eng et al., 2015). The gluetus medius is a hip abductor, and its anterior part is a hip flexor while its posterior part is an extensor (Beck et al., 2015). Thus, without sufficient measurements, it remains unclear how moments in the non-sagittal planes would change due to the motion in the sagittal plane.

Finally, to have a full picture of the joint positions well explored of their isometric capacities, we plotted a heat map of measurement frequency (Fig 12): Joint positions that frequently get measured in isometric experiments are plotted with larger and redder patches. It is evident that most of the measurements are for knee extension and ankle plantarflexion moments, and they are also limited to a few joint positions; e.g., knee moment at various knee angles with 90*^◦^* hip flexion, ankle moment at at various ankle angles with 0*^◦^* and 90*^◦^* hip flexion. Furthermore, to denote the importance of each joint position, we included the kinematics of incline walking, running, counter-movement jump, and loaded full squat, with indication of the moment required along the process. The rationale is that, the joint positions covered by these curves are *functional positions* that routinely occur in daily tasks, which we consider more superior in the joint space.

For accurate biomechanical analysis, moment measurements should be conducted in such functional positions rather than convenient positions in laboratory environments such as sitting and lying. However, it is rather clear that current measurements cover little of the region that these curves pass through. For example, hip flexion moment is mostly required between the mid-stance and mid-swing phases of running, where hip flexion is between −10*^◦^*–30*^◦^* and knee flexion between 15*^◦^*–120*^◦^*. In Fig 12 (top left, cyan curve), in the far left region of the plot where this part of the curve locates, only a few patches are lit up. This means that the available data are insufficient to support accurate biomechanical analysis for running, such as calibrating a musculoskeletal model to simulate a sprint. For more intensive motions such as jumping and squatting, there is a simultaneous increase of hip flexion, knee flexion, and angle dorsiflexion angles in the concentric phase and a simultaneous decrease in the eccentric phase, and the curves mainly travel through the diagonal direction of the plots. Nevertheless, the majority of this region has no measurement at all, and when measurements do exist, they are often from separate studies with potentially large variations in the data. Lacking constructive moment data in these function positions, it can be hard to deduce the moment–angle relation for further analysis. For instance, when squatting to the bottom position, where both hip flexion and knee flexion increase from 80*^◦^* to 110*^◦^*, is isometric knee extension moment maintained at some peak throughout the process? If so, is this peak different from that observed in Fig 11 (top right)? These questions are essential to estimating muscle activation in the knee, and they cannot be answered by the datasets at hand. Therefore, it is crucial to design experiments specifically covering the functional positions as indicated by Fig 12.

**Figure 11.**
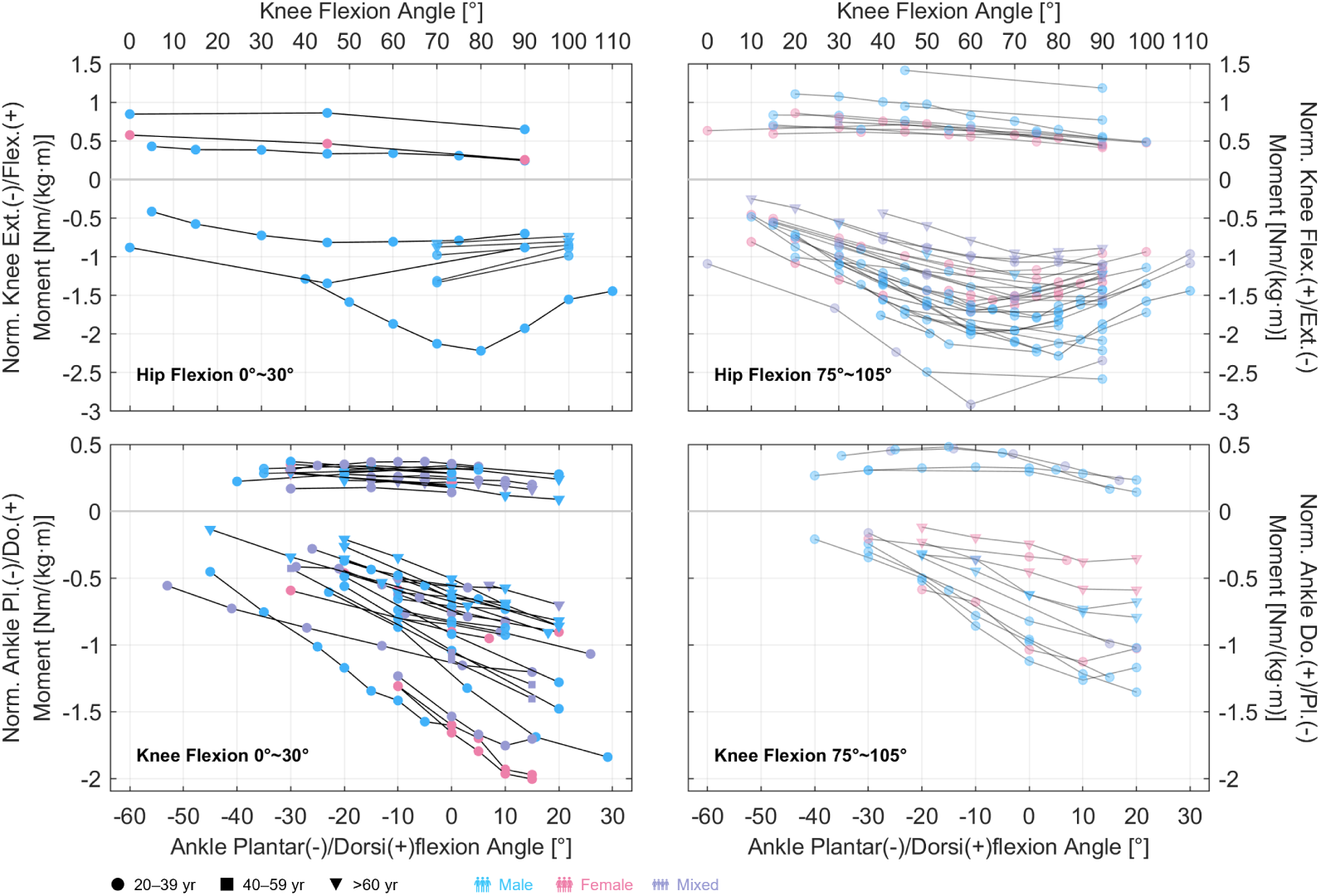
Relations of isometric knee extension/flexion and ankle plantar-/dorsiflexion moments with respective angles. Top left: Relation of knee moments when the hip is extended or slightly flexed. Top right: Relation of knee moments when the hip is highly flexed. Bottom left: Relation of ankle moments when the knee is extended or slightly flexed. Top right: Relation of ankle moments when the knee is highly flexed. Data are normalized by the product of subject height and weight.

**Figure 12.**
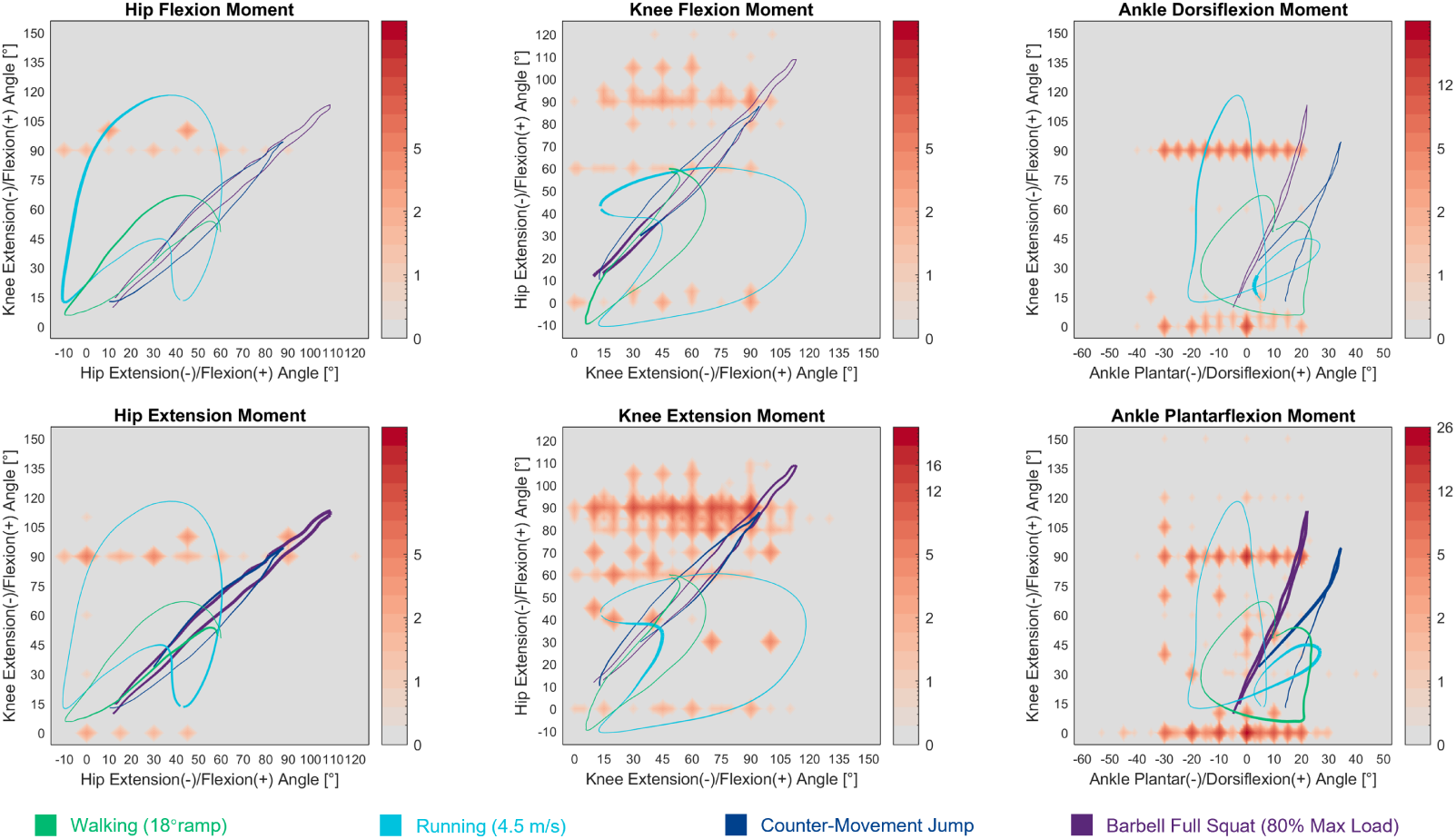
Measurement frequency of moments in the sagittal plane. The color assigned for each joint position denotes the number of studies that performed measurements at this position (color bar); e.g., 26 studies measured ankle plantarflexion moment at the position of 0*^◦^* ankle dorsiflexion and 0*^◦^* knee flexion. The colored curves denote the kinematics of dynamic motions, and the moment required at each position is indicated by the thickness of the segment.

### 4.4. Isokinetic Moment

A key point to notice in isokinetic moment datasets is how the data are obtained: As mentioned in the Methods section, they are different in whether the moment is the angle-specific, peak, or mean value during the tested RoM. Due to the moment–angle relation, the curves plotted based on these values would obviously be different in magnitude, but it remains unclear to many if measurements from different joint angles, when normalized against maximal isometric moment, would lead to the same moment–velocity relation. In other words, is the force–velocity relation length-dependent?

To answer this question, we may start from Hill’s (1938) famous experiment of shortening heat and dynamic constants, and see if his experiment falls into the category of length-specific, peak, or mean. Here, it must be noted that the relation Hill (1938) revealed is between constant load and mean velocity throughout an isotonic contraction, which is slightly different from the current muscle force–velocity relation. Also, there was no discussion over how muscle length would influence the results; in fact, based on Hill’s (1938) description, some of the tests clearly involved muscle contracting at non-optimal lengths. Nevertheless, as technology advances, it was demonstrated with force transducers that a similar relation holds for muscle force and instant velocity in both isotonic and isokinetic conditions (Gollapudi and Lin, 2009; Iwamoto et al., 1990; Mashima et al., 1972).

Importantly, it was further demonstrated that the force–velocity relation is length-dependent in many ways. Mashima et al. (1972) and Krylow and Sandercock (1997) showed that the maximal shortening velocity would decrease if the shortening starts from a length with smaller isometric force. This means the Hill constant *b*, hence the shape of the force–velocity curve, varies for different initial lengths; more specifically, the curve would appear steeper if the initial length is not optimal. The final length of contraction is also relevant, as muscle would experience force depression: In shortening contraction, muscle force is smaller than its maximal isometric value at the final length even when the velocity reaches zero, and the magnitude of decrease is greater for larger displacement (Herzog et al., 2000). Last but not least, Scott et al. (1996) showed that even in a single isokinetic contraction, regardless of concentric or eccentric, the extent by which force is decreased or increased would vary at different lengths.

Therefore, in isokinetic experiments, it is important that moment data are obtained at specific joint angles, rather than using the peak or mean values as a convenient replacement. It is equally critical to control for the tested RoM to eliminate the influence of initial isometric force and force depression. Nonetheless, this is often neglected, and many studies measured isokinetic moments merely as an indication of muscle strength for correlation analysis, so the majority of the datasets were recorded with peak values from different RoMs. This induces further variations in the isokinetic moment curves on top of the factors discussed in the Study Heterogeneity subsection.

In the main, the moment–velocity curves are rather chaotic (Figs 13–16). When normalized against measurements in the isometric condition, the relative moment curves present a pattern similar to the force–velocity relation, but the lower and upper bounds are noticeably different (Fig 17). In the concentric part of the force–velocity curve, muscle force would decrease to zero when shortening at a maximal velocity, while in the eccentric part, force would increase more than 1.6 times the isometric value (Krylow and Sandercock, 1997; Mashima et al., 1972). As shown in Fig 17, joint moment does not typically go below 25% in high concentric speeds, and rarely reaches over 125% in eccentric conditions. This is a major difference between the characteristics of muscle fiber and the muscle–tendon unit, and may be attributed to several reasons.

**Figure 13.**
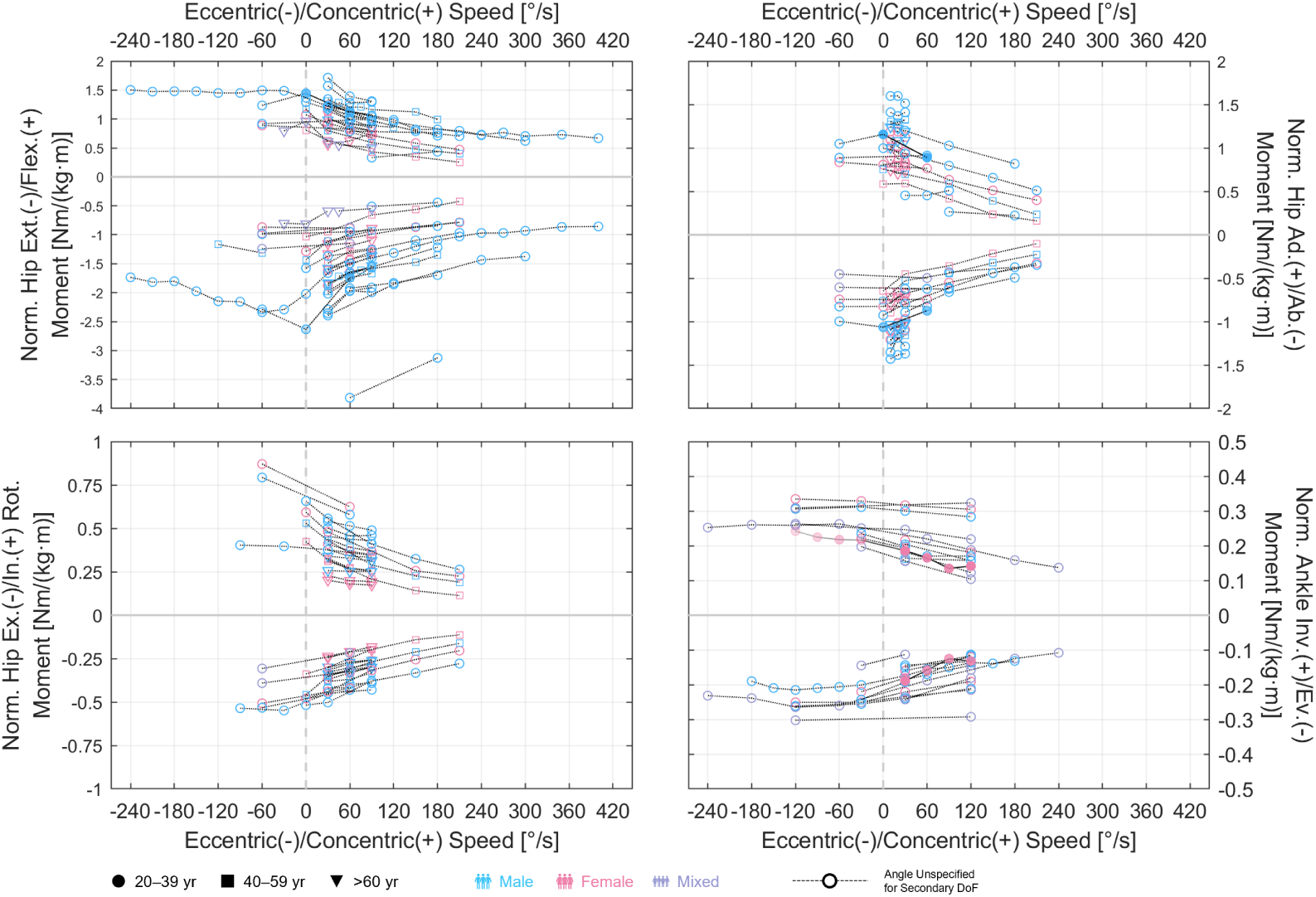
Relations of isokinetic moments with angular velocity in the hip and ankle. Top left: Hip extension/flexion. Top right: Hip ab-/adduction. Bottom left: Hip ex-/internal rotation. Bottom right: Ankle eversion/inversion. Data are normalized by the product of subject height and weight.

**Figure 14.**
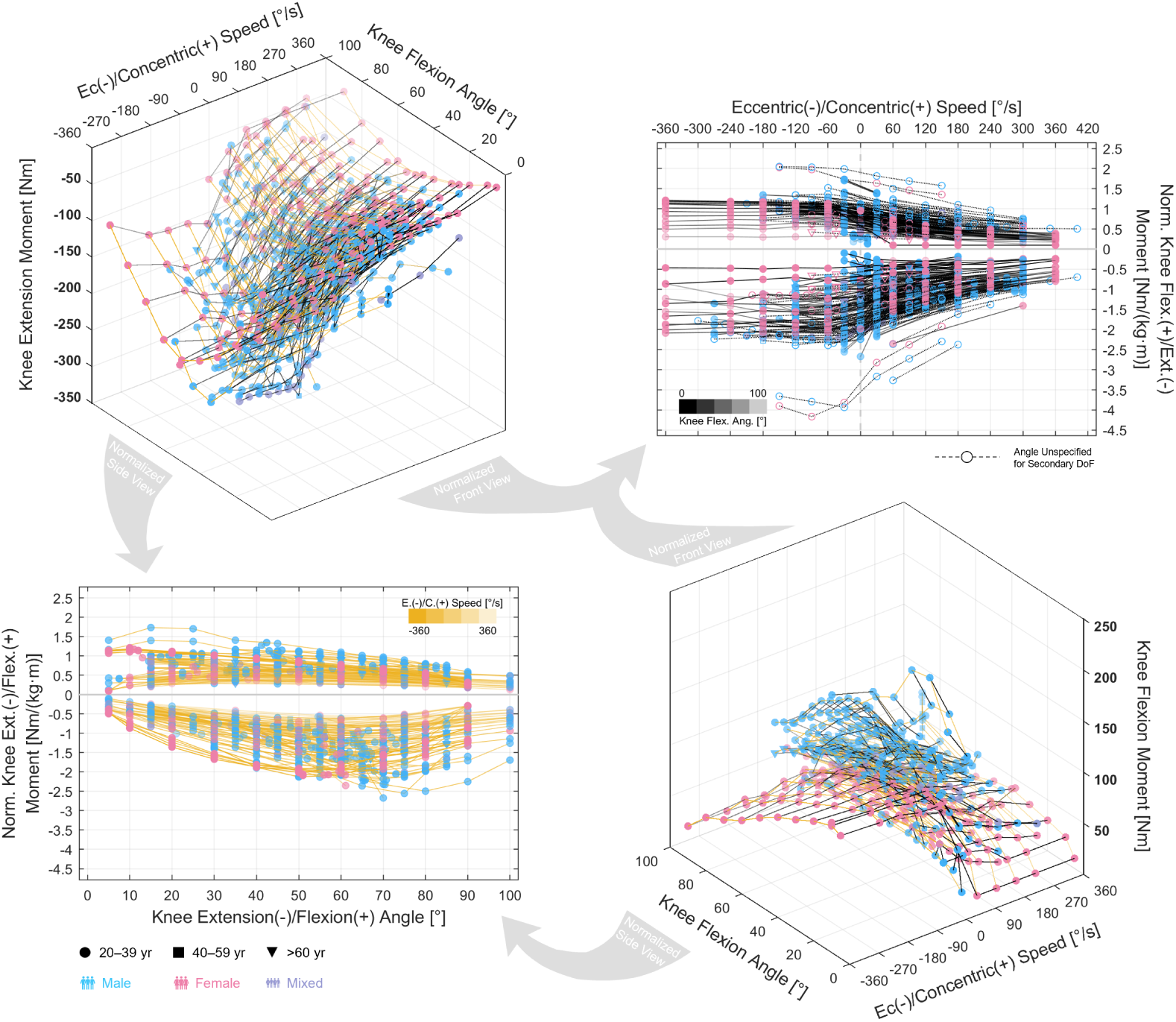
Relations of isokinetic knee extension/flexion moments. Top left: 3D relation of extension moment with angular velocity and angle. Top right: Normalized 2D relation with angular velocity. Bottom left: Normalized 2D relation with angle. Bottom right: 3D relation of flexion moment with angular velocity and angle. Data are normalized in the 2D plots by the product of subject height and weight. The transparency of the curve segment denotes the value in the relatively fixed DoF (color bar): with yellow for angular velocity and black for knee flexion angle.

**Figure 15.**
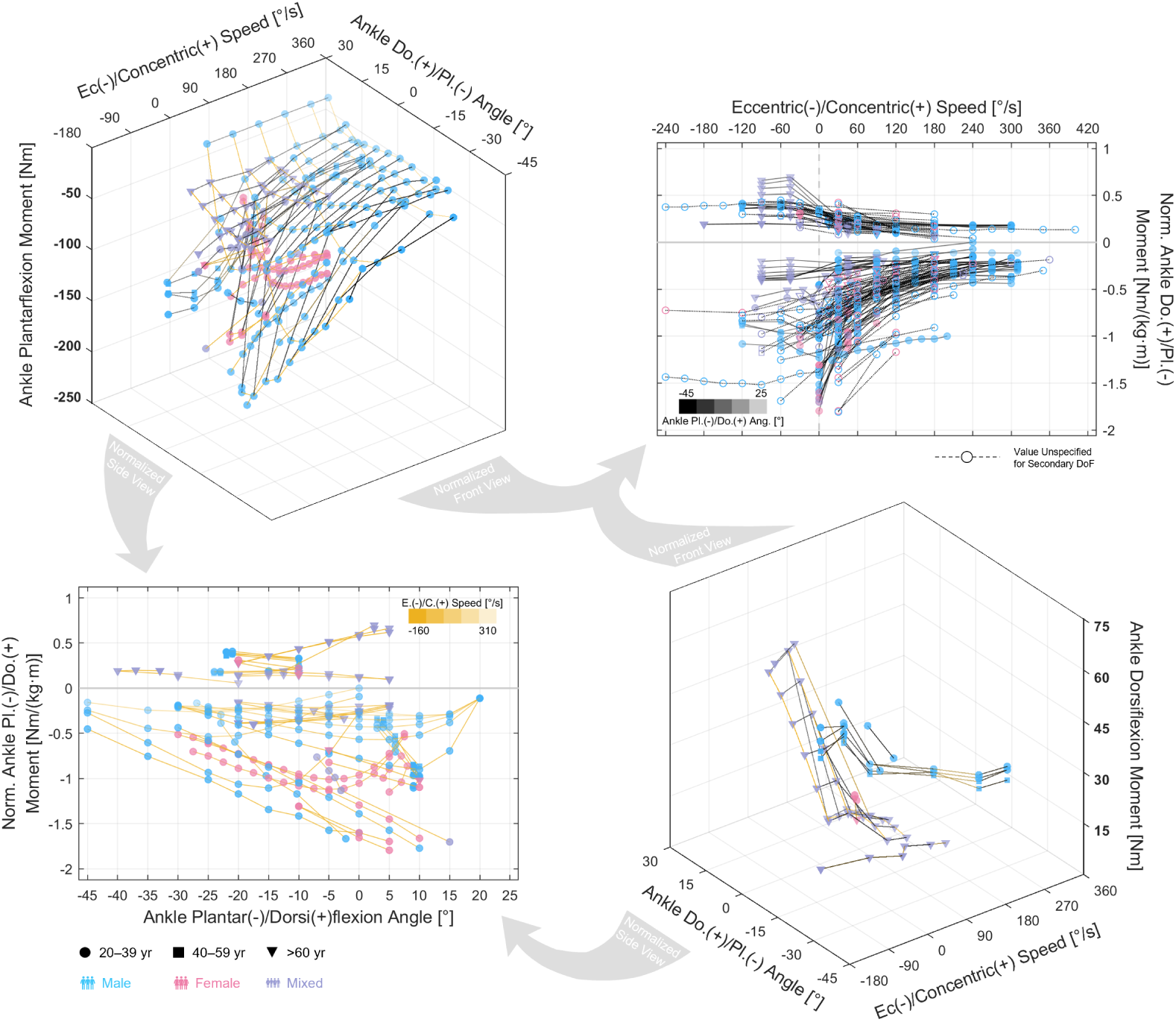
Relations of isokinetic ankle plantar-/dorsiflexion moments. Top left: 3D relation of plantarflexion moment with angle and angular velocity. Top right: Normalized 2D relation with angular velocity. Bottom left: Normalized 2D relation with angle. Bottom right: 3D relation of dorsiflexion moment with angular velocity and angle. Data are normalized in the 2D plots by the product of subject height and weight. The transparency of the curve segment denotes the value in the relatively fixed DoF (color bar): with yellow for angular velocity and black for ankle dorsiflexion angle.

**Figure 16.**
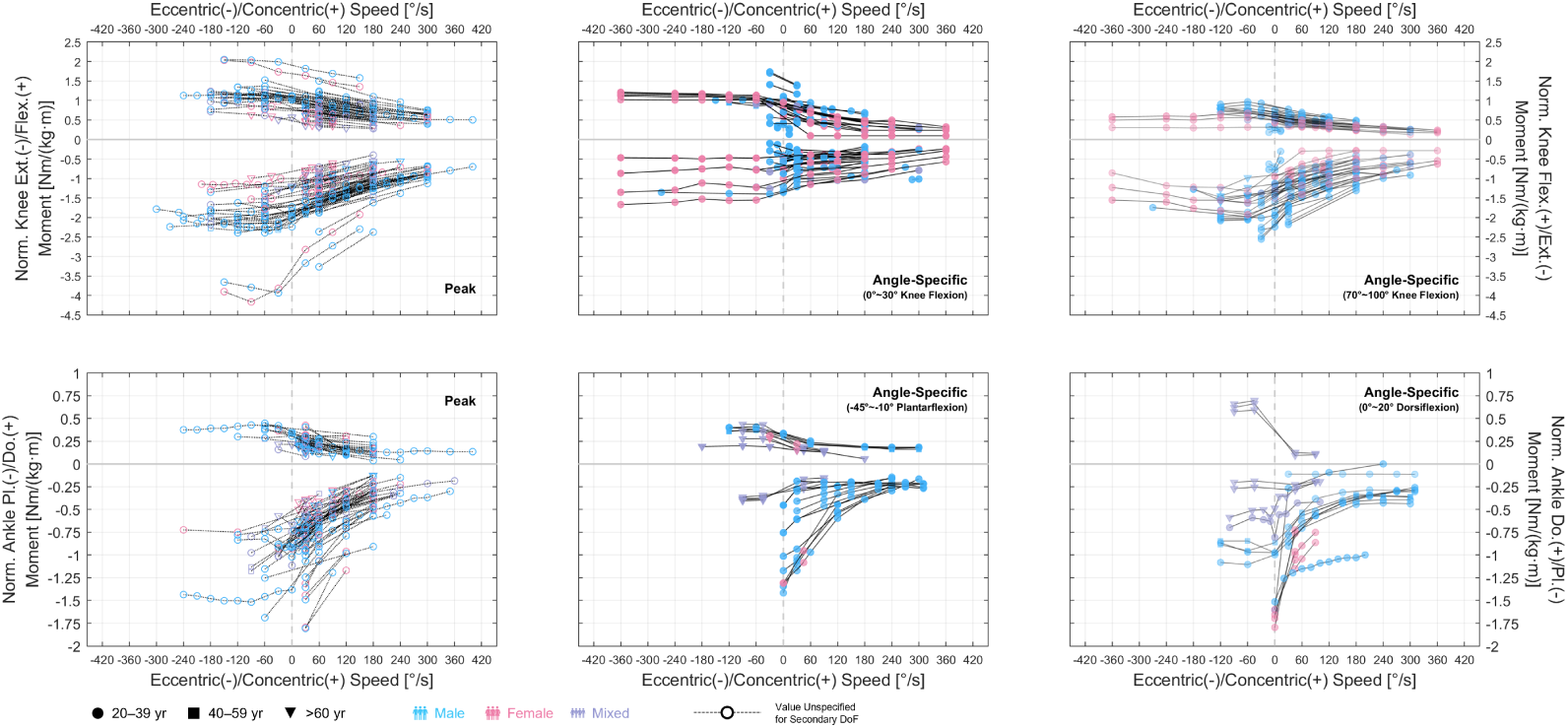
Relations of isokinetic knee extension/flexion and ankle plantar-/dorsiflexion moments with angular velocity. Top left: Relation of knee moments based on peak values. Top center: Relation of knee moments based on angle-specific values. Top right: Relation of knee moments based on angle-specific values from small knee flexion angles. Bottom left: Relation of ankle moments based on peak values from large knee flexion angles. Bottom center: Relation of ankle moments based on angle-specific values from ankle plantarflexion angles. Bottom right: Relation of ankle moments based on angle-specific values from zero or ankle dorsiflexion angles. Data are normalized by the product of subject height and weight.

**Figure 17.**
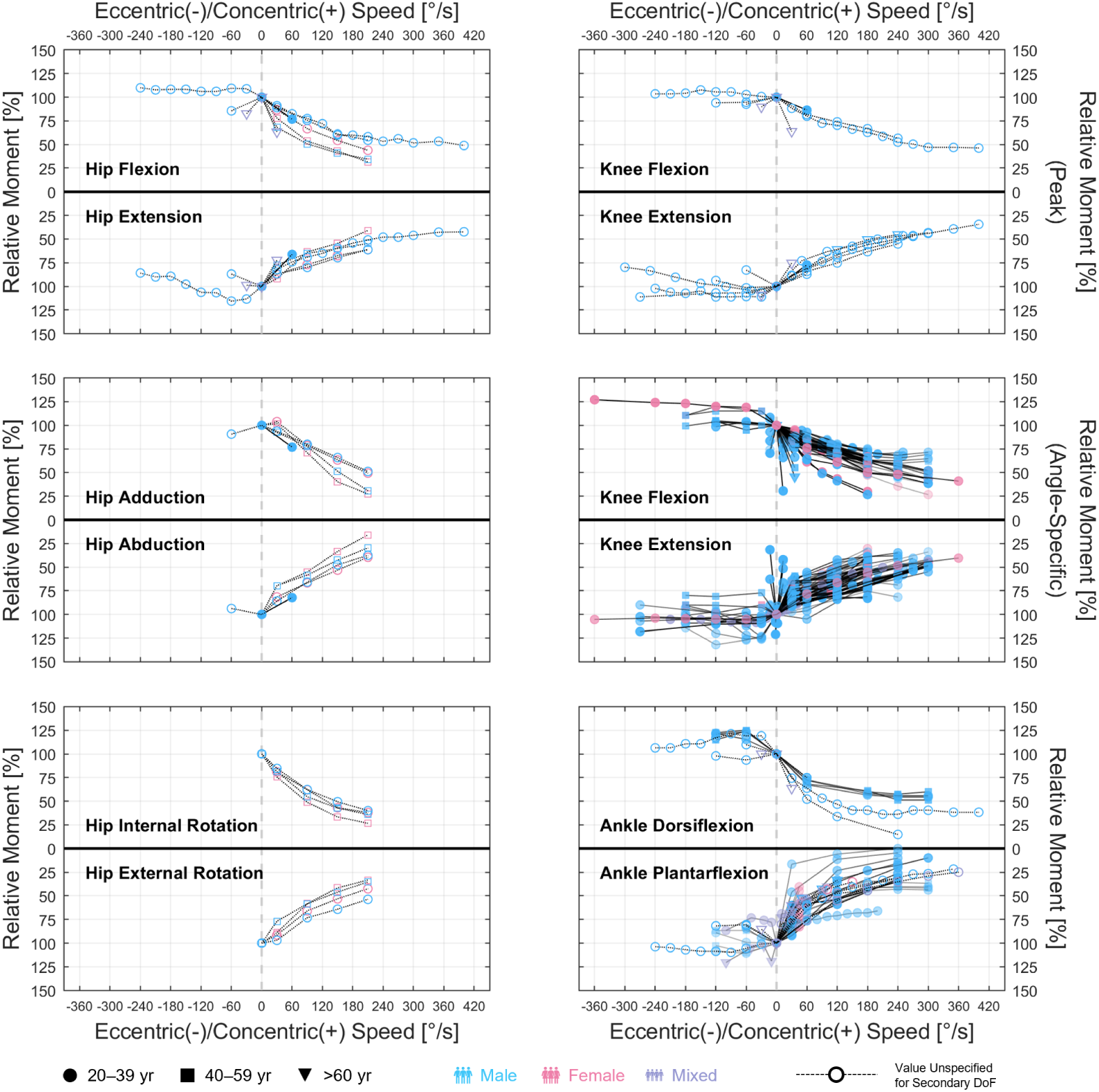
Relations of relative moments with angular velocity. Top left: Hip extension/flexion. Middle left: Hip ab-/adduction. Bottom left: Hip ex-/internal rotation. Top right: Knee extension/flexion, based on peak values. Middle right: Knee extension/flexion, based on angle-specific values. Bottom right: Ankle plantar-/dorsiflexion. Relative moment is calculated with the isometric value. For datasets based on peak values, the isometric value is only considered valid if it is the larger value of measurements obtained from at least two different angles. Otherwise, it is unclear whether the value is the peak isometric moment, and the increase will be overestimated in eccentric contractions while the decrease will be underestimated in concentric contractions.

First of all, the stretch of the elastic tendon during muscle contraction comprises up to 9.2% of muscle–tendon unit length change (Waugh et al., 2012), which decreases the shortening or lengthening velocity of muscle fascicles. For muscles with large pennation angles, such as the soleus, vastus medialis, and gluteus maximus (Ward et al., 2009), the velocity of muscle fascicles will be further decreased. The existence of connective tissues or the 3D architecture of the muscle may also cause a slow-down. Consequently, the fascicle might be contracting at a much slower speed than when calculated directly from joint rotational speed, and it may not operate on the far left and far right region of the force–velocity curve. Moreover, there could be neurological factors that prevent the exertion of maximal force in isokinetic conditions. Especially with eccentric contractions, subjects require training to get used to the experiment set-up, even still they might feel uncomfortable finishing the task. Then as aforementioned, the activation of agonists and antagonists are not necessarily full and null, and may even vary at different positions (Maganaris et al., 1998; Onishi et al., 2002; Salzman et al., 1993; Worrell et al., 2001), and so there is no reason to assume a constant and maximal muscle activation in an isokinetic contraction.

Despite the large variations in the isokinetic moment curves, readers are still encouraged to refer to them in biomechanical analysis. For example, in musculoskeletal modeling, the muscle is generally modeled with predefined force–velocity characteristics, and the resultant moment–angular velocity relation is not often validated. Suppose the joint moment ends up varying from 0%–150%, which is broader than the range suggested by the datasets, muscle activation would be underestimated in eccentric conditions and overestimated in concentric. Thus, it is necessary to validate the isokinetic moments in the model with measurements, in particular if the motions to simulate are highly dynamic.

### 4.5. Limitations

In order to gather as many relevant datasets as possible, we employed the common techniques in systematic review and worked our way from 8800 research results. Yet it would not be surprising if some of the featured studies are missed out. Joint moment is an ordinary type of data in biomechanical investigations, and its measurement is much more difficult to pin-point compared with the research questions typically addressed with meta-analysis. For this, we took a machine learning–like approach, where the keywords are tuned to ensure search accuracy in a calibration set. The format of the final keyword set is complicated but proves to be efficient: Of the 55 studies containing passive moment datasets, 40 appeared in searches based on keywords intended for the passive condition (73%). For the 117 isometric-related studies, 100 of them appeared in isometric-based searches (85%), while for the 163 isokinetic-related studies, the intended keywords led to 107 of them (66%). Meanwhile, we included searches for moment arm data using simpler keywords as a fail-safe for overfitting, where a study lacking one of the complex keywords fails to show up.

Even still, it is impossible to secure all relevant studies, simply because the field of biomechanics has thrived for decades with countless moment measurements. We are, however, confident that we have gathered the majority of datasets with quality results. When ranked with the frequency and place of appearing in searches, it is shown the bottom-ranked records are less likely to yield a target study: The target rate decreased from 14% in the first quartile of 4353 records to 9%, 6%, and 4% in the second, third, and fourth. Reasonably, we may deduce that no more than a few dozen studies would be added even with double the size of search results.

To accommodate potential amendments, we provide a log with details of the search workflow (Appendix A), where excluded records are labeled with the relevant data and the reason of exclusion. Readers may easily expand from the current results and gather datasets with other specific needs; e.g., moment in children or people with cerebral palsy. We believe this is by far the largest and most constructive collection of joint moment datasets, and with the advancement of artificial intelligence such as ChatGPT, we hope colleagues in the field will no longer need to manually repeat our work in the future.

Complete as it might be, our datasets do not cover joint moments in knee ab-/adduction, knee ex-/internal rotation, and ankle ex-/internal rotation, which we have predetermined to be insignificant motions. During our research, we have however identified a few studies measuring active moments in these DoFs with customized devices (Cammarata and Dhaher, 2008; Hester and Falkel, 1984; Zhang et al., 2001). Also, we did not discuss over the correlation of joint moment relations with sex, age or profession, which could be of interest to many. The metadata catalog we provide has a detailed record of subject information in each study, and readers may find many studies with a focus on these topics.

Finally, the joint moment relations we illustrate are based solely on datasets gathered from literature, and we did not conduct measurements of our own. Nevertheless, it is indispensable to have a general understanding of currently available datasets before a purposeful experiment can be designed. As shown in Fig 12, although there are 117 isometric datasets, they are crowded in positions less related to the kinematics of ordinary movements. With this in mind, we may then design experiments targeting important functional positions, and the results will be of more value for the biomechanics community.

## 5. Conclusion

A total of 985 datasets of passive, isometric, and isokinetic moment were collected from literature to illustrate joint moment–angle and moment– velocity relations in the hip, knee, and ankle. The overall pattern and magnitude, as well as the frequency of measurement are presented for 12 joint moments from six DoFs. The findings should provide valuable insight into joint kinetics, enhance biomechanical analysis, and improve the design of future experiments for moment measurement.

## 6. Acknowledgements

We thank Dr. Matthew Millard for the useful discussions. This work was supported by the Lighthouse Initiative Geriatronics by StMWi Bayern (Project X, grant no. 5140951).

## Appendix A. Supplementary material

Files associated with this article, including the search log and metadata catalog (.xlsx), datasets (.mat), and visualization scripts (.m), can be found online at *link-to-add*.

## Appendix B. Summary of studies and datasets

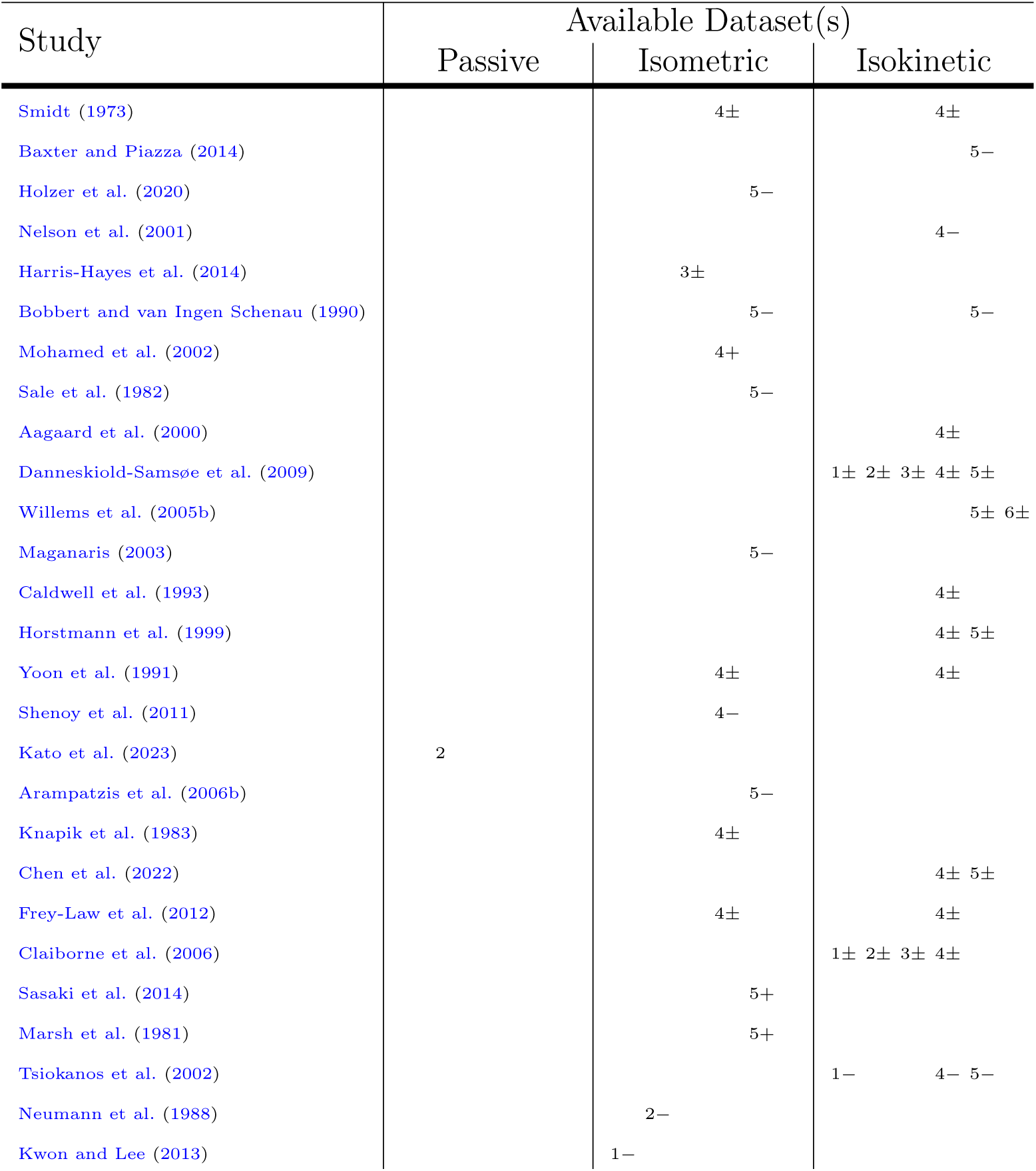
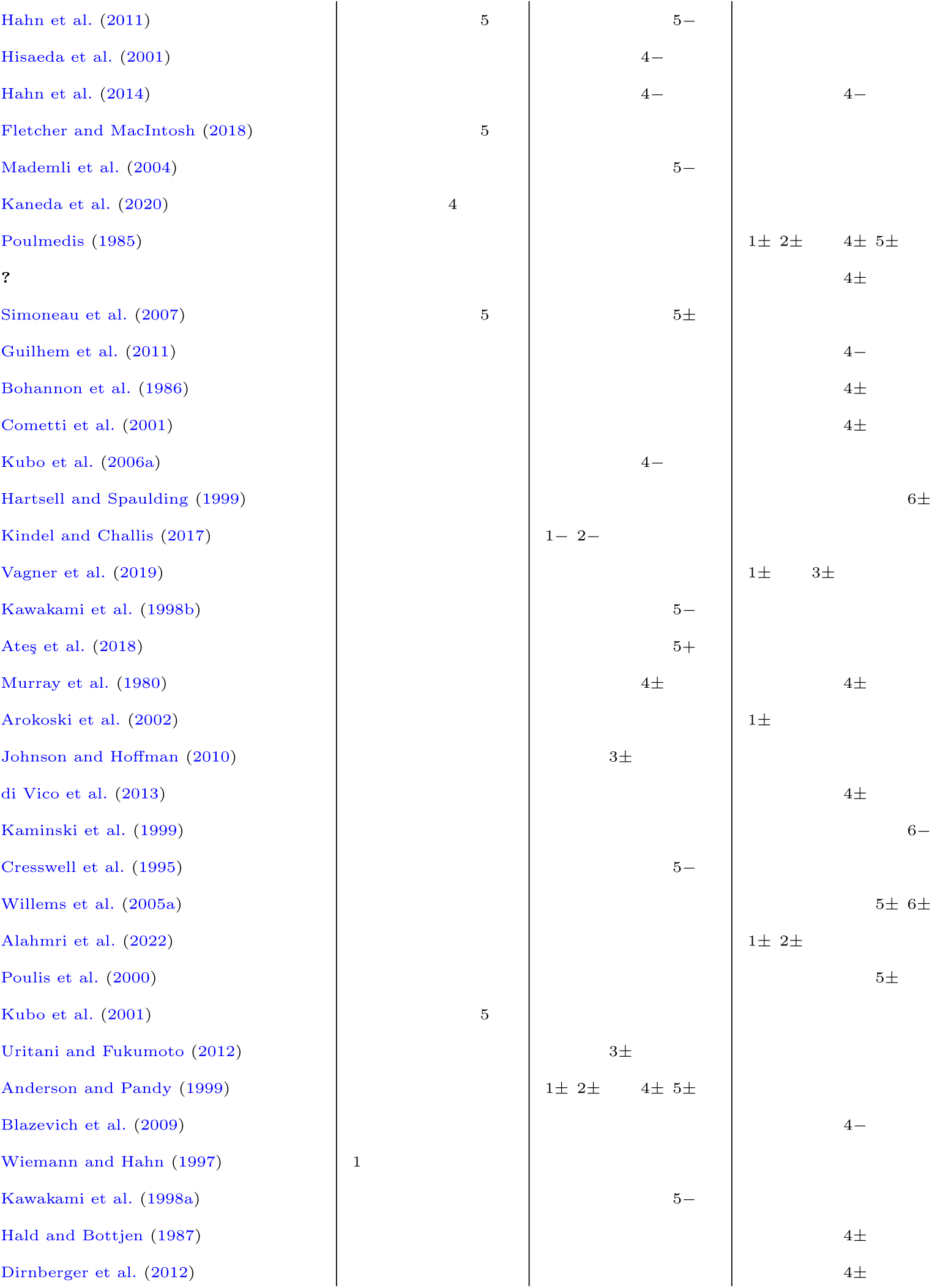
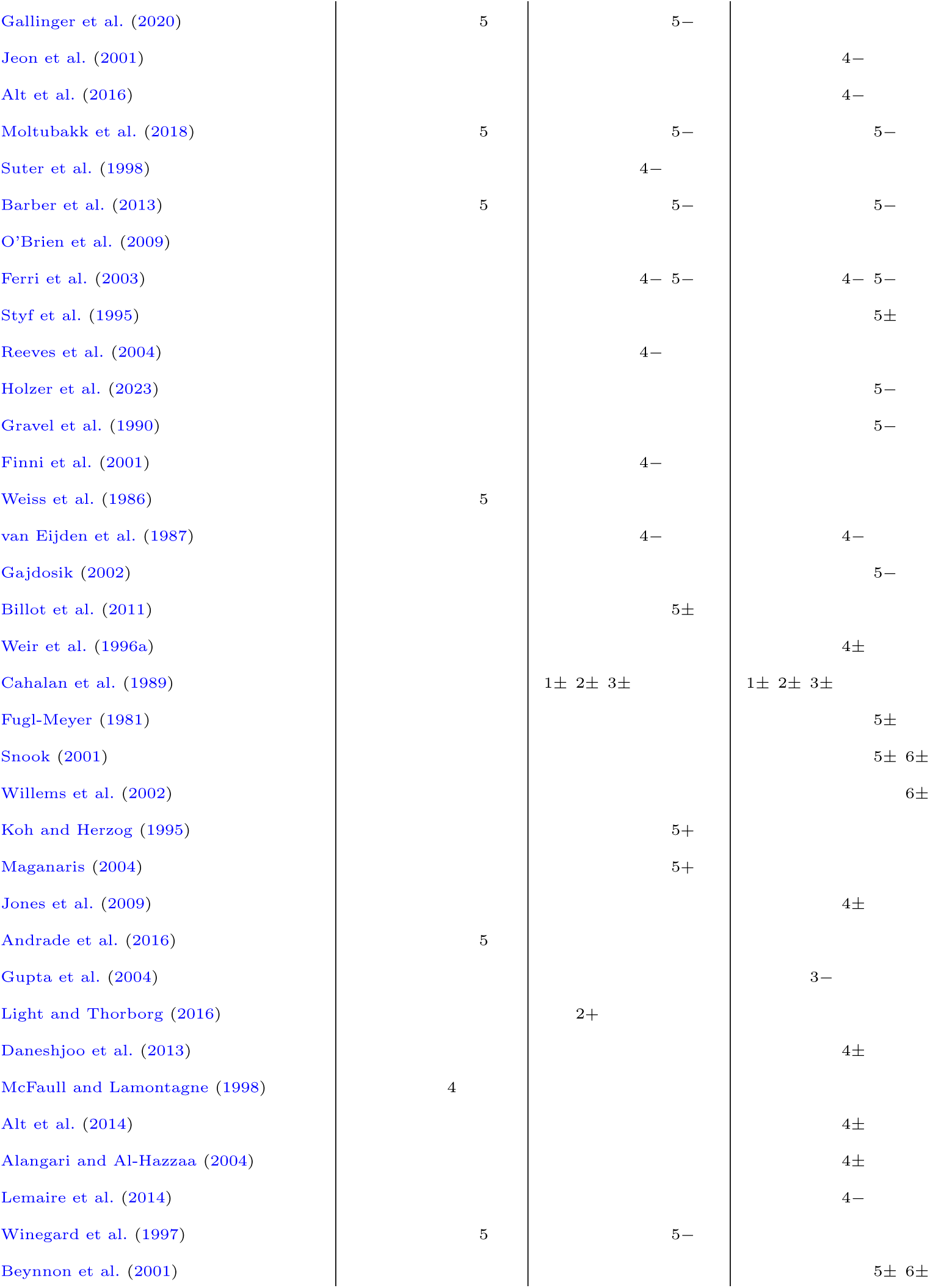
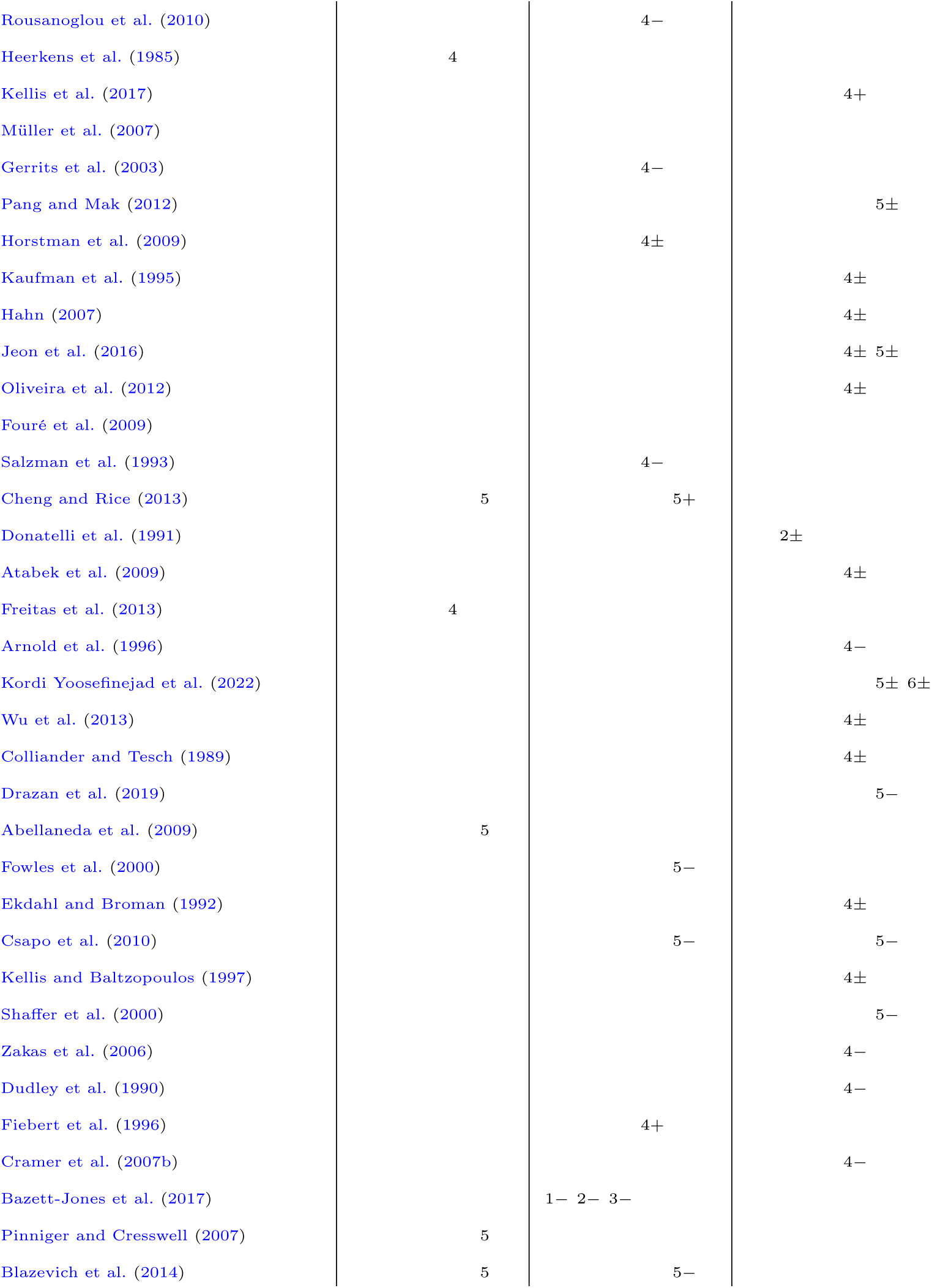
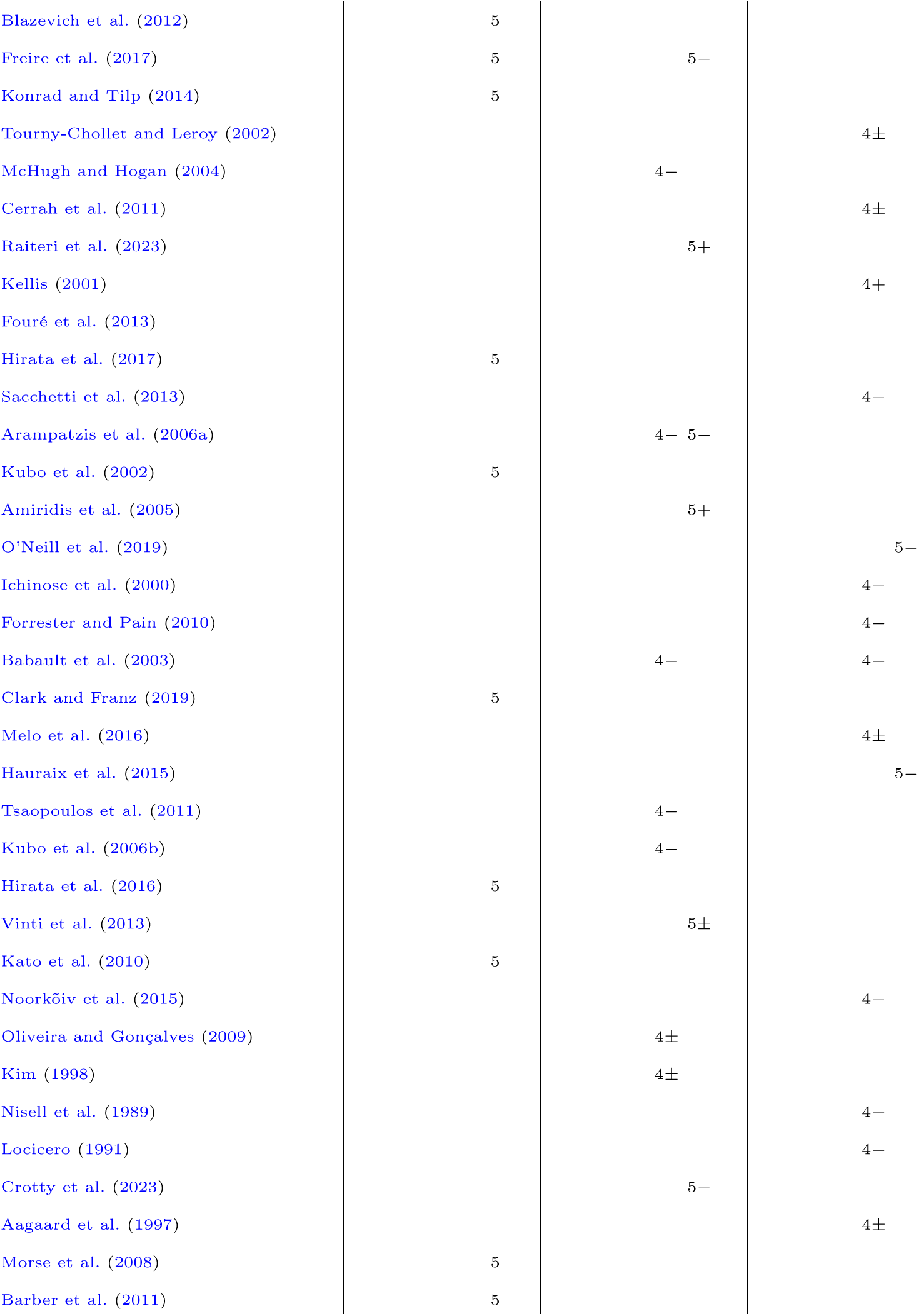
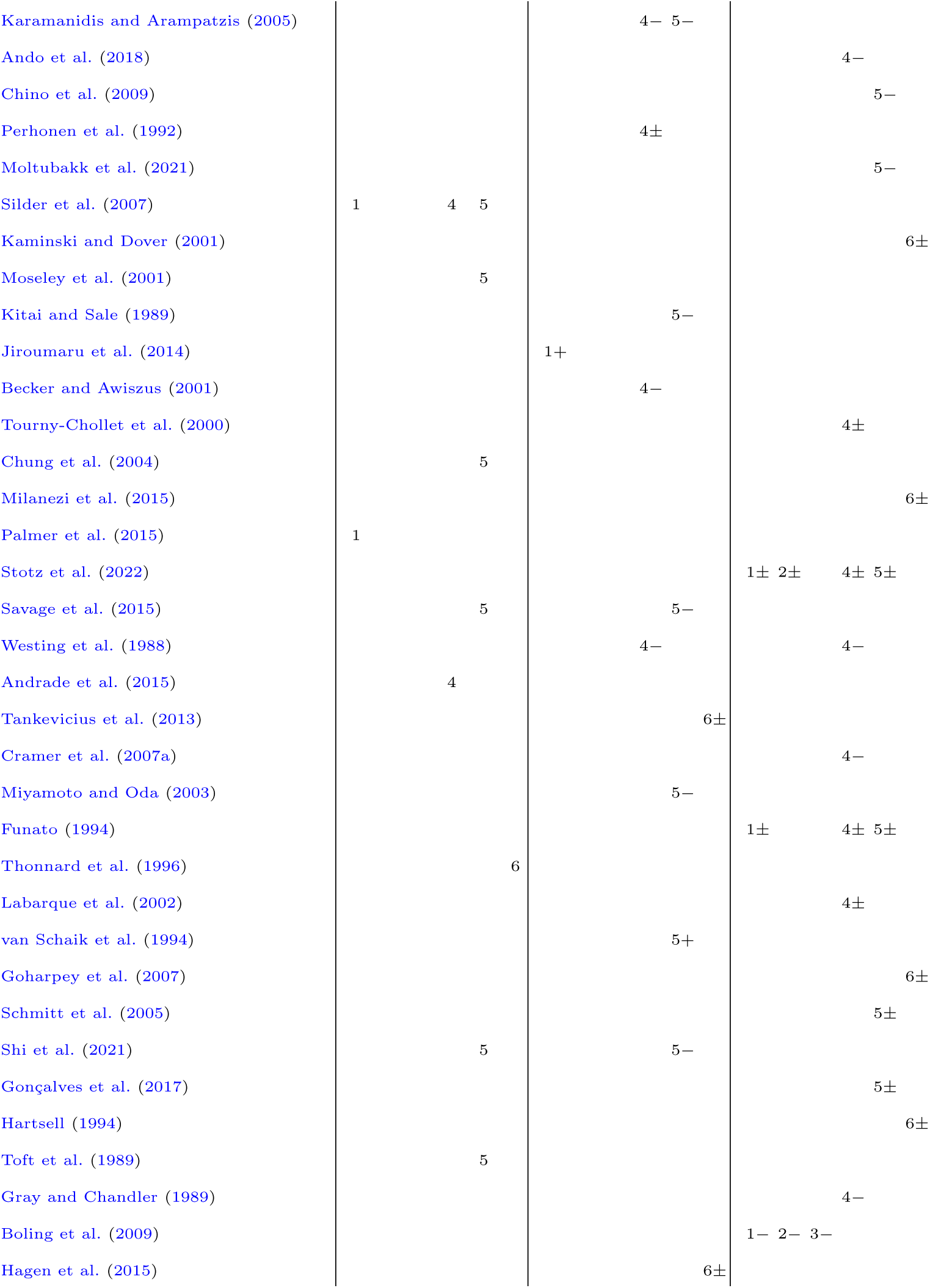
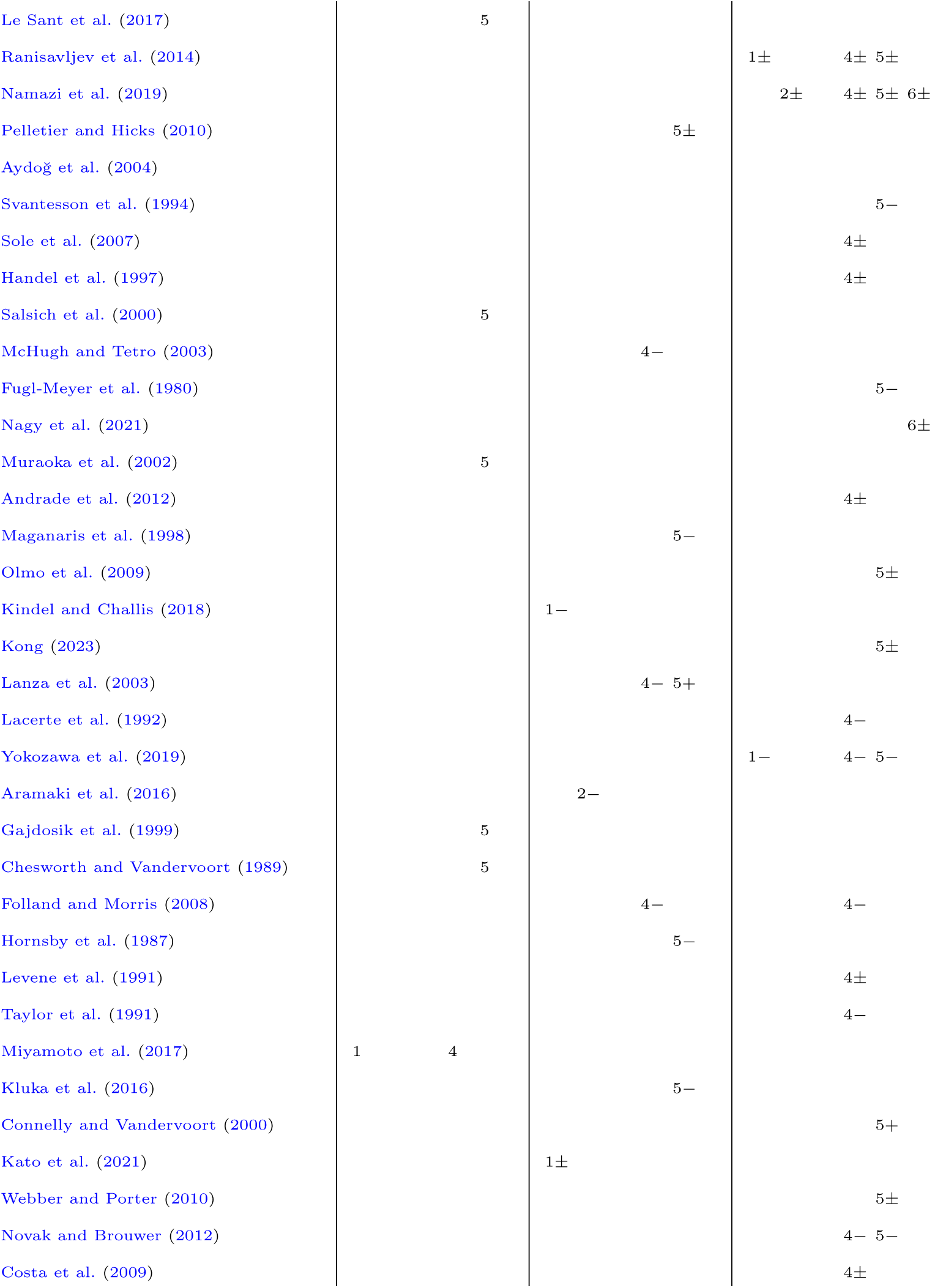
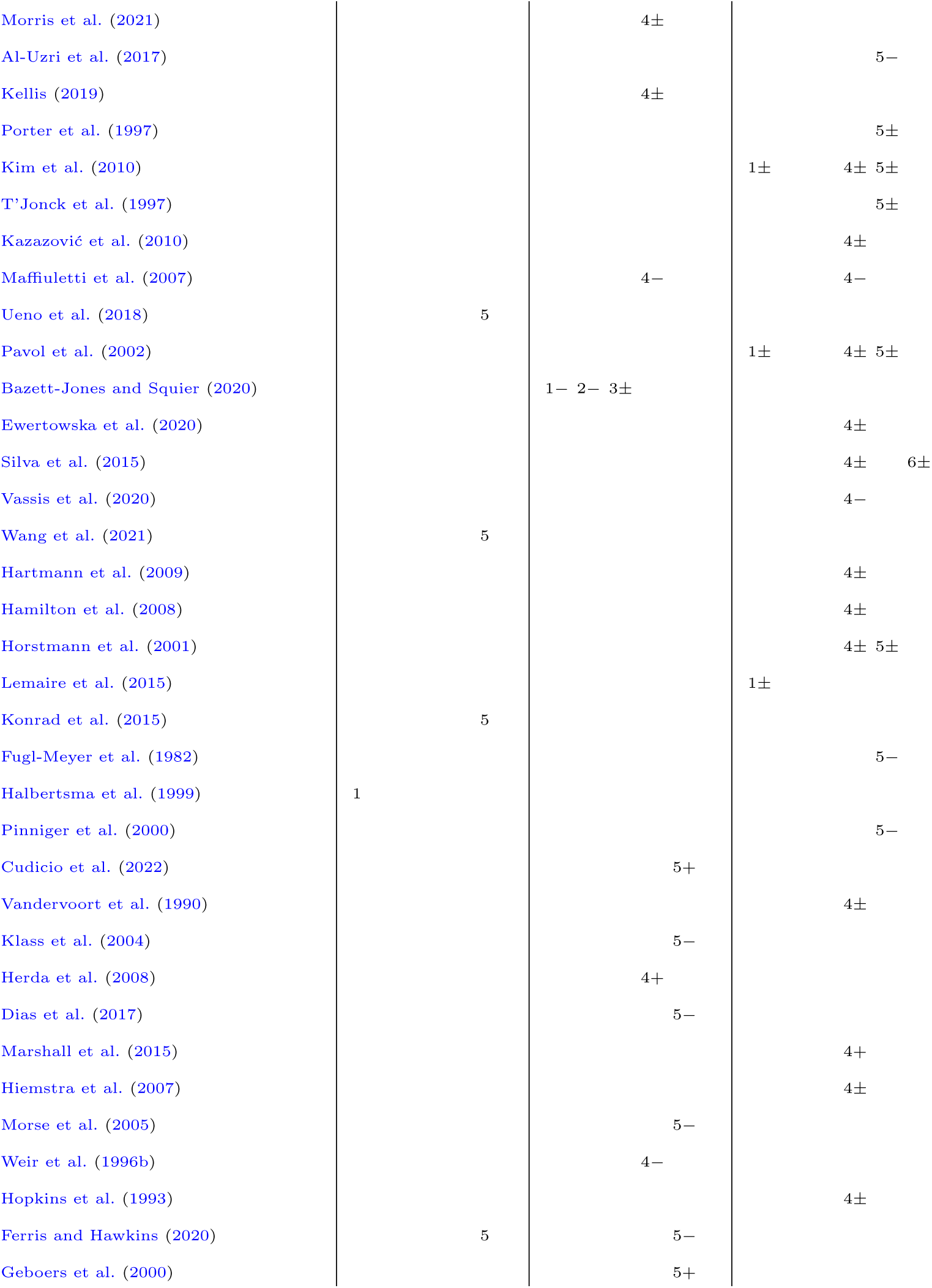
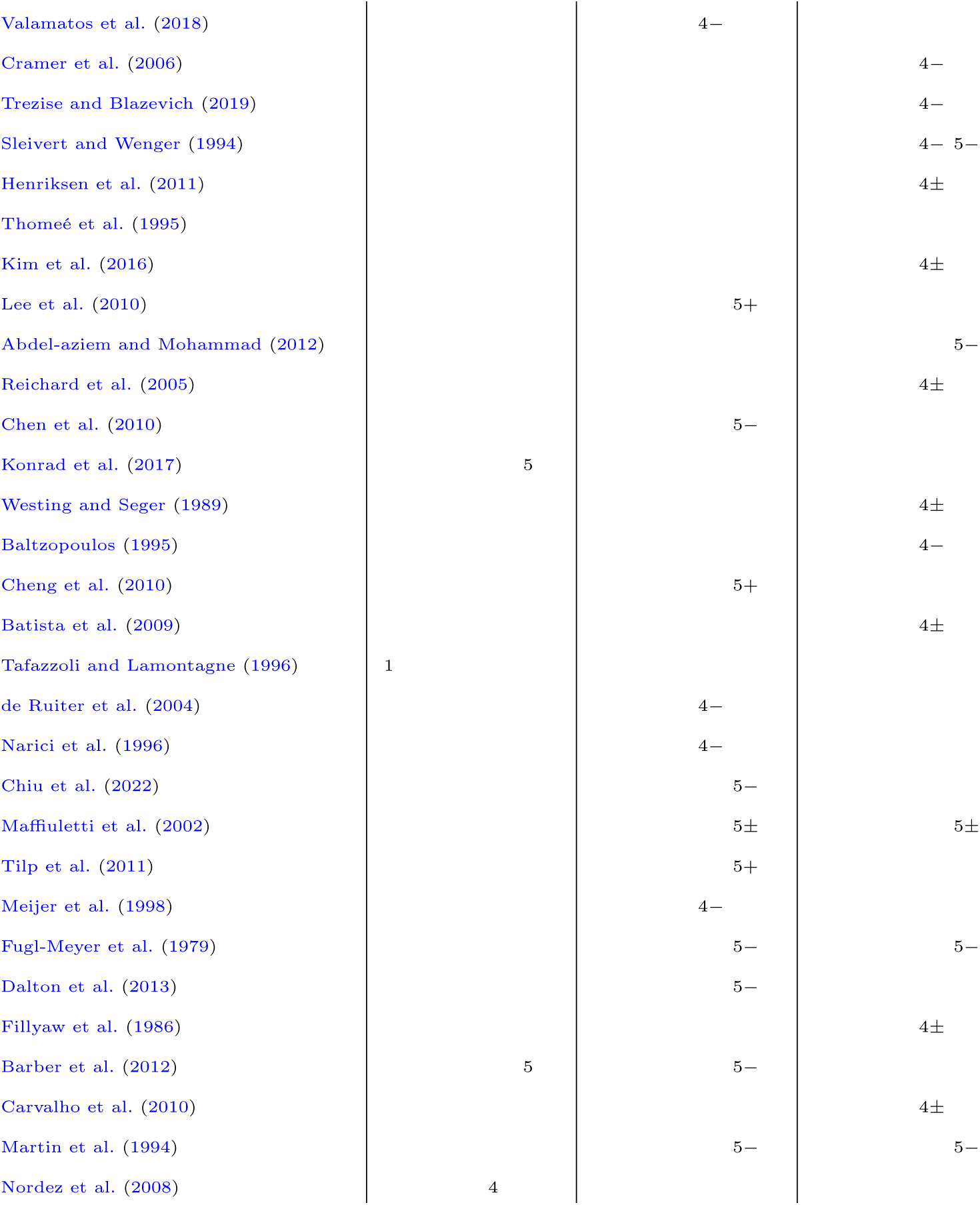

